# Long-PDE4D degradation attenuates left ventricular cardiomyocyte hypertrophic remodelling

**DOI:** 10.64898/2026.10.30.762532

**Authors:** Aleksandra Judina, Zhiyao Shi, Jiarong Fu, Ivan Sherstnev, Veronika Boichenko, Jiayue Ling, Madihah Hussain, Alina Zorn, Yuan Yan Sin, Houcheng Wang, Ao Li, Sasha Korogodski, Eduardo Torres, David Stirling, Kyle Chan, Cathy Swindlehurst, Iyobel Kibread, Iain Black, Stuart Nicklin, George S. Baillie, Julia Gorelik

**Author notes:** **Corresponding authors:** George S. Baillie, PhD, School of Cardiovascular and Metabolic Health, University of Glasgow, UK., Julia Gorelik, PhD, Imperial College London, 429, 4th Floor, ICTEM Building, Hammersmith Campus, United Kingdom. W12 0BZ.

## Abstract

Spatial compartmentalisation enables cyclic AMP (cAMP) to regulate distinct cellular functions, yet selectively targeting disease-associated signalling remains challenging. Phosphodiesterases constrain local cAMP responses, but conventional inhibitors offer limited isoform selectivity. Here we investigate whether targeted degradation of long phosphodiesterase-4D (PDE4D) isoforms can suppress pathological growth while preserving contractile function. Using adult rat and human ventricular cardiomyocytes and human myocardial slices, we show that the proteolysis-targeting chimera KTX-184 enhances compartmentalised cAMP signalling and attenuates hypertrophic remodelling. In rat cardiomyocytes, KTX-184 increases β-adrenergic cAMP responses in the cytosol and at the plasma membrane, including T-tubule and crest regions, and enhances nuclear protein kinase A activity. In contrast, phospholamban- and ryanodine receptor-associated cAMP nanodomains, calcium transients and sarcomere shortening remain unchanged. KTX-184 prevents angiotensin II-induced hypertrophic remodelling, preserves T-tubule organisation and normalises hypertrophic marker expression. In human cardiomyocytes, it increases cytosolic cAMP responses and prevents overload-induced hypertrophy in myocardial slices. These findings identify long PDE4D isoforms as targets for spatially selective modulation of cAMP signalling and support targeted degradation as an approach to suppress cardiac hypertrophy without increasing contractility.

## BACKGROUND

Cardiac hypertrophy is a progressive maladaptive response to sustained haemodynamic or neurohumoral stress that impairs ventricular relaxation, reduces pump efficiency, and predisposes to heart failure (HF), a condition affecting over 64 million people worldwide and carrying a five-year mortality exceeding 50%^1^. Current pharmacological standards of care, including β-blockers, ACE inhibitors, and calcium-channel blockers, improve symptoms and reduce hospitalisation but rarely reverse established hypertrophic remodelling, and patients with advanced disease continue to face substantial morbidity and mortality^1^. Therefore, there is an unmet need for therapies that directly target the molecular mechanisms driving pathological cardiac remodelling.

A central regulator of ventricular cardiomyocyte adaptation is the β-adrenergic receptor (β-AR)/cyclic adenosine monophosphate (cAMP)/protein kinase A (PKA) signalling axis. β-AR stimulation activates adenylyl cyclase, elevates intracellular cAMP, and engages PKA to phosphorylate targets that govern both contractile function and gene expression. Critically, this signalling is not uniform across the cell: β₁-ARs generate near-global cAMP elevations, whereas β₂-ARs produce spatially confined signals concentrated at T-tubule openings and caveolae^2–4^. These distinct cAMP nanodomains are actively maintained by phosphodiesterases (PDEs), which hydrolyse cAMP in close proximity to receptors and downstream effectors, thereby insulating spatially segregated signalling pools^5^. Disruption of nanodomain organisation, as occurs during hypertrophy and HF, redistributes β₂-AR signalling and alters transcriptional outputs, contributing to maladaptive remodelling^3,4^.

Among the PDE superfamily, the PDE4 subfamily plays a particularly prominent role in cardiomyocytes’ cAMP compartmentation. PDE4D, in particular, is expressed as multiple isoforms in ventricular myocytes, where long isoforms (e.g., PDE4D3, PDE4D5, PDE4D8, and PDE4D9) are tethered to distinct subcellular scaffolds: β-arrestin at β-ARs and muscle A-kinase anchoring protein β (mAKAPβ) at the nuclear envelope, thereby gating receptor-to-nucleus PKA signalling ^6–8^. Structurally, long PDE4D isoforms differ from short isoforms by containing both upstream conserved region 1 (UCR1) and UCR2 domains, which mediate homodimerisation, AKAP-scaffold integration, and PKA-dependent feedback activation via UCR1 phosphorylation^9,10^. Short isoforms, which lack UCR1, exist as monomers and are excluded from these regulatory molecular complexes^11^. Importantly, only long PDE4D isoforms have been detected in ventricular cardiomyocytes^12–16^, providing the structural basis for an isoform-selective therapeutic strategy.

Despite the clear rationale for targeting PDE4D in cardiac disease, pharmacological approaches have been limited by poor selectivity. Broad PDE3 inhibition, explored to restore impaired β-AR responsiveness in HF, increased arrhythmia risk and mortality in clinical trials, precluding its chronic use^17^. PDE4-selective inhibitors demonstrate anti-inflammatory and cardioprotective properties and, importantly, lack direct positive inotropy ^18–21^. However, first-generation PDE4 inhibitors were not designed with isoform-or nanodomain-selectivity in mind, contributing to systemic side effects including emesis and gastrointestinal intolerance, and inconsistent cardiac efficacy^8^. The ability to selectively remove a specific PDE4D isoform from a defined subcellular compartment, without globally suppressing PDE4 catalytic activity, would represent a conceptually distinct and potentially superior therapeutic approach.

Proteolysis-targeting chimeras (PROTACs) offer precisely this capability through an “event-driven” pharmacological mechanism. Rather than occupying an active site, PROTACs are bifunctional molecules that simultaneously engage a target protein and an E3 ubiquitin ligase, inducing proximity-dependent ubiquitination and subsequent proteasomal degradation of the target^22^. Because each PDE4D isoform carries a unique N-terminal extension that dictates its subcellular localisation and protein-protein interactions, these regions present tractable handles for isoform-selective degradation. KTX-184 is a novel PROTAC built on an apremilast-derived PDE4-binding warhead linked to a cereblon-recruiting ligand; cereblon serves as the substrate receptor of the CRL4 E3 ubiquitin ligase complex. Notably, a prior PDE4 PROTAC using a related apremilast scaffold selectively degraded only short-form PDE4D^23^, which is not appreciably expressed in cardiac tissue, highlighting the importance of linker and warhead engineering in achieving the desired isoform selectivity.

Here, we demonstrate that KTX-184 selectively degrades long PDE4D isoforms in adult left ventricular cardiomyocytes, relieving sarcolemmal and nuclear cAMP constraints, enhancing β-AR-to-nucleus PKA signalling, and attenuating hypertrophic remodelling, without perturbing RyR2- or phospholamban (PLN)-centred contractile nanodomains or altering calcium (Ca^2+^) transients. Using complementary approaches including FRET-based reporters of compartmentalised cAMP and PKA activity, scanning ion conductance microscopy (SICM)-FRET, high-throughput contractility assays, an angiotensin II (AngII)-induced hypertrophy model, and human myocardial tissue slices, we established targeted PDE4D degradation as a precision strategy for anti-hypertrophic therapy.

## METHODS

### Chemicals

All reagents were acquired from Sigma (UK) unless otherwise specified. KTX-184 and apremilast were obtained from Katalytic Therapeutics, Inc. (San Diego, CA, USA). Stock solutions of KTX-184 or apremilast were prepared in dimethyl sulfoxide (DMSO) and diluted into experimental buffers immediately before use. Final DMSO concentrations did not exceed 0.1% (v/v) in any experiment.

### Computational model of PDE4D, cereblon and KTX-184 interactions

A structural model of the ternary complex formed by PDE4D, KTX-184, and cereblon was generated using Molecular Operating Environment (MOE) software (version 2024.0601; Chemical Computing Group, Montreal, Canada). Crystal structures of PDE4D bound to apremilast (PDB: 7CBQ) and cereblon bound to thalidomide (PDB: 4CI1) were used as templates. Structures were prepared by removing non-essential subunits, metals, and water molecules, followed by energy minimisation using the MOE QuickPrep tool. The PDE4D-binding and cereblon-binding moieties of KTX-184 were docked independently into their respective binding pockets using the co-crystallised ligands as templates. Docking poses were manually inspected, and the optimal binding mode was selected for ternary complex assembly using MOE Method4B. The resulting complex ensemble was clustered and ranked by binding energy to identify the best-scoring pose.

### Ethics statement

All animal experiments were conducted in accordance with the UK Animals (Scientific Procedures) Act 1986, the European Directive 2010/63/EU, and institutional guidelines at Imperial College London. All procedures were approved by the Imperial College London Animal Welfare and Ethical Review Body.

Human tissue studies were conducted in accordance with the Human Tissue Act 2004 under the Imperial College Healthcare NHS Trust Tissue Bank licence. Human donor hearts were obtained through NHS Blood and Transplant (NRES ethics reference: 16/LO/1568; study numbers 67 and 106). Written informed consent was obtained from all donors or their next of kin.

### Isolation and culture of adult rat left ventricular cardiomyocytes

Adult left ventricular (LV) cardiomyocytes were isolated from male Sprague-Dawley rats (250-350 g; Charles River, UK) by Langendorff perfusion and enzymatic digestion at 37 °C, as previously described ^24,25^. Briefly, digested LV tissue was mechanically disaggregated and filtered through a cell strainer to obtain single-cell suspensions. Cardiomyocytes were plated on laminin-coated 25 mm borosilicate glass coverslips (VWR, USA) and maintained in Minimum Essential Medium (MEM; Gibco, UK) supplemented with 1% antibiotic-antimycotic (Sigma-Aldrich, USA), 1% L-glutamine, and 9 mmol/L NaHCO₃, at 37 °C and 5% CO₂ for 48-h. For the *in vitro* hypertrophy model, isolated rat LV cardiomyocytes were treated with angiotensin II (AngII; 20μmol/L; Abcam, UK) for 24-h, as previously described ^26–29^. Where indicated, KTX-184 (1nmol/L) was co-administered with AngII.

### Human LV cardiomyocyte isolation and culture

Human LV cardiomyocytes were isolated from donor heart LV tissues by enzymatic digestion as previously described ^30,31^. Tissue sample were maintained in ice-cold low-Ca^2+^ solution (120mmol/L NaCl, 5.4mmol/L KCl, 5mmol/L MgSO_4_, 5mmol/L Pyruvate, 20mmol/L glucose, 20mmol/L taurine, 10mmol/L HEPES, 5mmol/L NTA, 1 μmol/L CaCl_2_; pH = 7.2-7.4,), minced into ∼1mm^3^ pieces, and incubated in protease XXIV solution (0.36 mg/mL in 120mmol/L NaCl, 5.4mmol/L KCl, 5mmol/L MgSO_4_, 5mmol/L pyruvate, 20mmol/L glucose, 20mmol/L taurine, 10mmol/L HEPES, 150μmol/L CaCl_2_) for 45 min at 37 °C with continuous oxygenation, followed by collagenase Type V (1mg/mL) for 15 min. Tissue was mechanically disaggregated, filtered, and centrifuged at 1,000 rpm for 1 min. The cardiomyocyte pellet was resuspended in fresh MEM media for subsequent experiments.

### H9c2 cardiomyoblast culture and pharmacological treatments

H9c2 rat cardiomyoblasts were kindly supplied by the Fuller laboratory and were originally obtained from the European Collection of Authenticated Cell Cultures (ECACC). Cells were maintained in Dulbecco’s modified Eagle’s medium (DMEM) supplemented with 10% foetal bovine serum and 1% penicillin-streptomycin at 37°C in a humidified atmosphere containing 5% CO₂. Unless otherwise specified, cells were harvested 24-h after drug treatment.

To assess concentration-dependent effects on PDE4 protein abundance, cells were treated with KTX-184 at 1, 10 and 100 nmol/L and 1 and 10 μmol/L for analysis of PDE4D long forms, or at 10 and 100 nmol/L and 1 and 10 μmol/L for analysis of PDE4B long forms. Apremilast (100 nmol/L) and vehicle (0.1% DMSO) served as the respective comparator conditions for these experiments. For time-course experiments, cells were treated with KTX-184 (100 nmol/L) for 1, 2, 4, 8 or 24 h; apremilast (100 nmol/L, 24 h) served as the comparator.

To assess whether KTX-184-induced loss of PDE4D long forms required proteasome activity, cells were treated with KTX-184 or apremilast in the presence or absence of bortezomib (100 nmol/L). A bortezomib-only condition was also included. Following treatment, cell lysates were analysed by immunoblotting for PDE4D or PDE4B long forms, with GAPDH used as the loading control. Recombinant PDE4D5 and PDE4B3 standards were included in the corresponding concentration-response experiments. Protein abundance was quantified by densitometry and expressed relative to the designated comparator condition. In the proteasome-inhibition experiments, lysates were additionally analysed by anti-ubiquitin immunoblotting to assess the accumulation of ubiquitinated proteins.

### Immunocytochemistry and live-cell membrane staining of rat LV cardiomyocytes

For immunocytochemistryfluorescence, rat LV cardiomyocytes were fixed, permeabilised, and incubated overnight at 4°C with primary antibodies against PDE4D (rabbit anti-PDE4D, 1:100; Proteintech 12918-1-AP), PDE4B (1:100; Invitrogen 40-1400), or *α*-actinin (1:100; Sigma A7811) in blocking buffer (5% BSA, 0.2% Tween-20, and 5% normal goat serum). Cells were then incubated with Alexa Fluor 488-or Alexa Fluor 546-conjugated secondary antibodies and mounted onto glass coverslips using ProLong Gold with DAPI (Thermo Fisher Scientific).

For live-cell T-tubule membrane staining, cells were incubated with 10 µmol/L Di-8-ANEPPS (Thermo Fisher Scientific) in Ca^2+^- and Mg^2+^-free HBSS for 3 min at room temperature, 24-h after drug treatments^32^.

Fluorescence images were acquired using a Zeiss LSM710 confocal microscope. T-tubule staining density was quantified using Fiji (ImageJ v2.3.0), and regularity was assessed by Fourier transform-based power spectral analysis in MATLAB, as described previously^33^.

### Immunofluorescences staining of H9c2 cell line

H9C2 cells were plated on an 8-well chamber slide and left for at least 24-h for the cell to attach. Cells were treated for 24-h before cell culture medium was removed to stop treatment and cells were fixed with 4% paraformaldehyde at room temperature for 10 min. Cells were washed thrice with filtered PBS and then incubated with WGA-488 (fluorescent membrane stain) for 10 min at 37°C as per manufacturer’s instructions. Cells were then permeabilised in 0.1% Triton X-100 in PBS for 7 min. Cells were again washed thrice with PBS before being blocked with 10% donkey serum and 2% BSA in PBS for 1-h at room temperature. Following this, cells were washed thrice with PBS before being incubated with primary antibodies (diluted in 5% donkey serum and 1%BSA in PBS) in a DIY humidifier chamber and left at 4°C overnight.

The next day, cells were washed with PBS thrice for 5 min with gentle agitation on a shaker before being incubated with the appropriate secondary antibodies (diluted in 5% donkey serum and 1%BSA in PBS) for 1 hour at room temperature. Following this, cells were washed with PBS thrice with gentle agitation. After the final wash, slides were dehydrated and mounted onto coverslips with ProLong™ Gold Antifade Mounting Solution with DAPI (Invitrogen, #P36931). Slides were left to dry in the dark for at least 24 hours before visualising. The primary antibodies used were PDE4D (In-house, 1:4000) and secondary antibody used was Donkey anti-Sheep 594 (Alexa Fluor #A11016; 1:500)

The immunostaining was visualised using a Zeiss LSM880 laser-scanning confocal microscope with oil immersion objective (63×/1.4 NA plan apochromat lens) and analysed using ImageJ (NIH, Bethesda, MA, USA).

### Neonatal rat cardiomyocyte isolation, culture and hypertrophy assessment

Neonatal rat cardiomyocytes were isolated by enzymatic digestion (Miltenyi Biotec) and mechanical dissociation using a gentleMACS dissociator. Cell suspensions were filtered through a 70-μm strainer, centrifuged at 1,000 × g for 5 min and preplated for 1 h to enrich cardiomyocytes. Non-adherent cells were collected and seeded onto laminin-coated coverslips in supplemented M199 medium.

Cells were pretreated with KTX-184 or apremilast (both 100 nmol/L) for 24 h before stimulation with angiotensin II (500 nmol/L) for a further 48 h. Control cultures received vehicle (0.1% DMSO) or angiotensin II alone. Cells were fixed and stained with FITC-conjugated phalloidin (1:100 in 1% BSA) and incubated for 1-h to visualise F-actin. Cell area was quantified from fluorescence images using Zeiss LSM880 laser-scanning confocal microscope with oil immersion objective (63×/1.4 NA plan apochromat lens) and analysed using ImageJ.

### Adenoviral FRET biosensor transduction and FRET microscopy

Adenoviral vector encoding FRET biosensors were amplified in HEK-293-AV cells and purified using the ViraBind^TM^ Adenovirus Miniprep Kit (Cambridge Bioscience) per the manufacturer’s instructions. The following biosensors were used: AKAR3-NES (cytosolic PKA activity reporters based on PKA-dependent phosphorylation of a 14-3-3τ substrate sequence) and AKAR3-NLS (nuclear PKA activity) ^34,35^; Epac1-camps (cytosolic cAMP)^36^; pmEpac2 (plasma membrane-localised cAMP) ^37^; RII_Epac (PKA RII-associated cAMP) ^6,38^; EPAC1-JNC (junctional cAMP) ^39,40^; and EPAC1-PLN (phospholamban-associated cAMP) ^41^.

Isolated LV cardiomyocytes were transduced at 100 MOI in MEM (1% antibiotic-antimycotic; 37 °C, 5% CO₂). At 24-h post-transduction, cells were treated with KTX-184, apremilast, or DMSO vehicle. FRET measurements were performed 48-h post-transduction. For concentration-response experiments, adult rat LV cardiomyocytes expressing Epac1-camps were treated with KTX-184 (10 pmol/L, 100 pmol/L, 1 nmol/L, 10 nmol/L, or 100 nmol/L) or DMSO vehicle for 24 h before FRET measurements. To assess the dependence of KTX-184-mediated cAMP responses on proteasome activity, Epac1-camps-expressing adult rat LV cardiomyocytes were treated with KTX-184 (1 nmol/L) in the presence or absence of bortezomib (100 nmol/L), with bortezomib alone and DMSO vehicle included as controls. Cells were stimulated with isoprenaline at biosensor-appropriate concentrations: 3 nmol/L for AKAR3-NLS ^35^ and AKAR3-NES ^34^, 10 nmol/L for pmEpac2 ^37^, EPAC1-JNC ^39,40^, EPAC1-PLN ^41^, and RII_Epac ^6,38,42^; or 100 nmol/L for Epac1-camps^36^. The KTX-184 concentration–response and proteasome-inhibition experiments both used isoprenaline at 100 nmol/L. Responses were normalised to a saturating cocktail of forskolin (10 μmol/L) and 3-isobutyl-1-methylxanthine (IBMX; 100 μmol/L) ^3^. For subtype-selective β-AR stimulation, ICI 118,551 (50 nmol/L; β_2_-antagonist) or CGP-20712A (100nmol/L; β_1_-AR antagonist) were applied together with isoprenaline solutions ^3^. FRET images were captured every 6 sec, and CFP/Venus emission ratios quantified using ratiometric multi-cell FRET microscopy ^43^.

### High-throughput sarcomere contractility and Ca^2+^ transient measurement

Sarcomere shortening and intracellular Ca^2+^ transients were measured using a CytoCypher high-throughput system (IonOptix, Westwood, MA, USA). Cardiomyocytes were loaded with Fura-2 AM (1 μmol/L) for 45 min, followed by a 15 min wash in Hank’s balanced salt solution (HBSS; Sigma), as previously described^30,44^. Cells were field-stimulated at 1 Hz (20mV), and sarcomere length and Fura-2 fluorescence ratio were recorded using the CytoSolver Transient Analysis Tool (IonOptix). Baseline measurements were followed by acute stimulation with ISO (1 μmol/L)^25^. The proportion of arrhythmic events was quantified over a 10 sec recording window. Resting cardiomyocyte morphology was assessed from brightfield thumbnail images to extract two-dimensional morphological parameters of cardiomyocytes^30,45^.

### Scanning Ion Conductance Microscopy with FRET imaging

Rat LV cardiomyocytes transduced with pmEpac2 or AKAR3-NLS were treated with KTX-184 (1 nmol/L), apremilast (1 nmol/L), or DMSO (0.1%) 24-h post-transduction. Nanoscale topographical maps (10 x 10 µm) of the cardiomyocytes’ surface were obtained using hopping-probe Scanning Ion Conductance Microscopy (SICM)^46^. ISO (10 µmol/L) was locally delivered via the SICM nanopipette by voltage application either at a T-tubule opening or a crest membrane region^4,38^. Simultaneous CFP and Venus fluorescence images were acquired every 6 sec using a digital ORCA-Flash 4.0 camera (Hamamatsu Photonics, UK) controlled by Micro-Manager 1.4 (open source, Vale Lab, University of California, USA) integrated with ImageJ. The FRET ratio was calculated as the Venus / CFP pixel intensity ratio minus a bleed-through correction factor of 0.956^38^.

### RNA isolation and RT-qPCR

Total RNA was extracted from adult rat LV cardiomyocytes using QIAzol Lysis Reagent™ (QIAGEN, 79306) per the manufacturer’s protocol. RNA concentration and purity were assessed by NanoDrop™ 2000c spectrophotometry (Thermo Scientific); samples with A_260_/A_280_ and A_260_/A_230_ ratios ≥ 1.8 were used for downstream analysis.

For mRNA quantification, cDNA was synthesised using the GoScript™ Reverse Transcriptase kit (Promega, A5001) and RT-qPCR performed with the GoTaq® qPCR Master Mix (Promega, A6001) on a QuantStudio™ 6 Flex system (Thermo Scientific). For miRNA quantification, cDNA was synthesised using the TaqMan™ MicroRNA Reverse Transcription Kit (Applied Biosystems™, 4366596) and amplified with the TaqMan™ Fast Advanced Master Mix (Applied Biosystems™, 4444557). All reactions were run in triplicate, and relative expression was calculated using the 2^(-ΔΔCt) method, with *Ubc* as the mRNA and *U6* as the miRNA reference gene. Primer sequences are provided in Supplementary Tables 1 and 2.

### Human myocardial donor tissue slices preparation and culture

Human tissue slices were prepared as previously described^47^. Full-thickness tissue blocks (∼1.5 cm^2^) from the donor LV free wall were mounted epicardial-side down and sectioned into 300 μm myocardial slices using a high-precision vibratome in oxygenated, ice-cold Tyrode’s solution containing 2,3-butanedione monoxime (BDM). Slices were trimmed to uniformly aligned regions and mounted on custom stretchers with biocompatible end attachments. Sarcomere length was set to 2.2μm (physiological preload; control) 2.4μm (mechanical overload). Slices were cultured for 72-h at 37 °C in Medium 199 supplemented with insulin-transferrin-selenium (ITS), ascorbic acid, and penicillin/streptomycin, with continuous oxygenation and medium recirculation^47^. For the AngII hypertrophic model, slices at physiological preload were treated with AngII (100 μmol/L) for 72h. The anti-hypertrophic effect of KTX-184 (100 nmol/L) was assessed in over-load slices.

### WGA staining of human donor tissue sections

Tissue slice sections were washed in PBS (5 min), fixed in 4% paraformaldehyde (10 min, room temperature), washed three times in PBS, and permeabilised with 0.25% Triton X-100 in PBS (15 min). After blocking (1-h, room temperature) in 1% BSA, 5% normal goat serum, and 0.25% Triton X-100. Sections were incubated with Alexa Fluor 647-conjugated WGA (1:200) and Hoechst (1:1,000) for 2-h at room temperature. Sections were washed twice in PBS, mounted in Citifluor, and cover-slipped.

### Western blotting of rat LV cardiomyocytes

Rat LV cardiomyocytes were lysed in RIPA buffer supplemented with phosphatase inhibitors (PhosSTOP, Roche), protease inhibitors (cOmplete Mini, Roche), and 1 mmol/L dithiothreitol (DTT). Protein concentration was determined by bicinchoninic acid (BCA) assay (Thermo Fisher Scientific). Equal protein amounts were combined with LDS sample buffer, denatured at 95 °C for 10 min, resolved on 4%-12% Bis-Tris SDS-PAGE gels, and transferred to nitrocellulose membranes. Membranes were blocked in 5% phosphoblocker solution in TBS-T (0.1% Tween 20) for 1-h at room temperature, then incubated overnight at 4 °C with previously validated in-house sheep anti-PDE4D or sheep anti-PDE4D5 primary antibodies^48^. After three washes in TBS-T, membranes were incubated with the appropriate IRDye - conjugated secondary antibodies (1:10,000; LI-COR Biosciences) for 1-h at room temperature. Fluorescent signals were detected and quantified using a LI-COR Odyssey CLx imaging system. For accessing the phosphoralyted protein, blots were blocking in 5% phospho-blocker (Cell Biolab, AKR-103) in TBS-T for 1-h at room temperature, the phospho blots were incubated with various phospho-antibodies and the total antibodies as controls in 1% phospho-blocker in TBS-T. The following antibodies were used for this study: p-THI (1:1000, Cell signalling), cTHI (1:1000, Abcam), p-PLB (1:1000, Millpore), PLB (1:1000, Thermofisher), p-HSP20, (1:3000, Abcam), HSP20 (1:3000,Upstate).

### Western Blotting of H9c2 cell line

H9c2 cellular lysates were prepared in lysis buffer (25 mmol/L Tris/HCl pH 7.4, 150 mm NaCl, 1% NP-40, 1 mmol/L EDTA, 5% glycerol) with complete EDTA-free protease inhibitor cocktail tablets (Roche, West Sussex, UK). Proteins were separated by SDS/PAGE (4–12% Bis-Tris gels) at 150V for 15 minutes, which was followed by an increase to 200V for 45 minutes. Gels were transferred onto nitrocellulose membranes at 30V for 90 minutes. The membranes were blocked in 0.5x Intercept (TBS) Blocking Buffer (LI-COR) for 1 h at room temperature, followed by incubation with primary antibody dilution in Intercept T20 (TBS) Antibody Diluent overnight (LI-COR) at 4 °C with gentle shaking. Primary antibodies used were sheep anti-PDE4D (1 in 1000; in house), anti-PDE4B (1 in 1000; in house), mouse anti-GAPDH (# 60004-1-Ig, 1:80 000; Proteintech), rabbit anti-GAPDH (#ab181602, 1:5000; Abcam), and mouse anti-ubiquitin (#sc-8017, 1:1000; Santa Cruz). Membranes were washed with TBST and then incubated with fluorescently labelled secondary antibodies (1: 15 000) for 1 h at room temperature. The secondary antibodies used were Donkey anti-mouse were IRDye 680 (#926-68072), IRDye 800 Goat anti-mouse 800 (#926-32210), IRDye 680 Donkey anti-goat, (#926-68074), IRDye Donkey anti-goat (#926-32214), IRDye 680 Goat anti-rabbit Goat (#926-68071) and IRDye 800 Goat anti-rabbit (#926-32211) from LI-COR Biosceinces (Lincoln, Nebraska, USA). Membranes were washed three times for 10 min each in 1× TBST in between antibodies and before visualisation. Images were acquired using the Li-Cor Odyssey CLx Imaging System. All densitometry analyses were performed using LI-COR Image Studio and normalised to GAPDH. Representative images are shown in grayscale.

### Statistics

All statistical analyses were performed using GraphPad Prism 10 Software (v.10.5.0). Data are presented as mean ± Standard Error of the Mean (S.E.M.). Group comparisons were made using an unpaired two-tailed *t*-test or Mann-Whitney test; one-way ANOVAs with Tukey’s post hoc multiple comparison test; or Kruskal-Wallis tests; or two-way ANOVA with Šídák’s multiple comparison test, as appropriate. A *P*-value <0.05 was considered statistically significant.

## RESULTS

### KTX-184 induces targeted degradation of PDE4D, but not PDE4B, in cardiomyocytes

To investigate the role of long PDE4D degradation in cardiomyocytes, we selected KTX-184, a novel long PDE4D-degrading molecule, based on prior evidence supporting an anti-hypertrophic and therapeutic role for PDE4D inhibition in cardiomyocytes ^20,21^.

Chemically, KTX-184 consists of PDE4, and E3 ligase-binding moieties connected via an optimised linker^48^ (Figure 1a, b). The PDE4D-binding moiety of KTX-184 was derived from the apremilast scaffold, an orally available PDE4 inhibitor approved for the treatment of psoriatic arthritis^49^. Computational modelling confirmed that KTX-184 can simultaneously engage PDE4D and cereblon into a ternary complex (Figure 1a).

**Figure 1.**
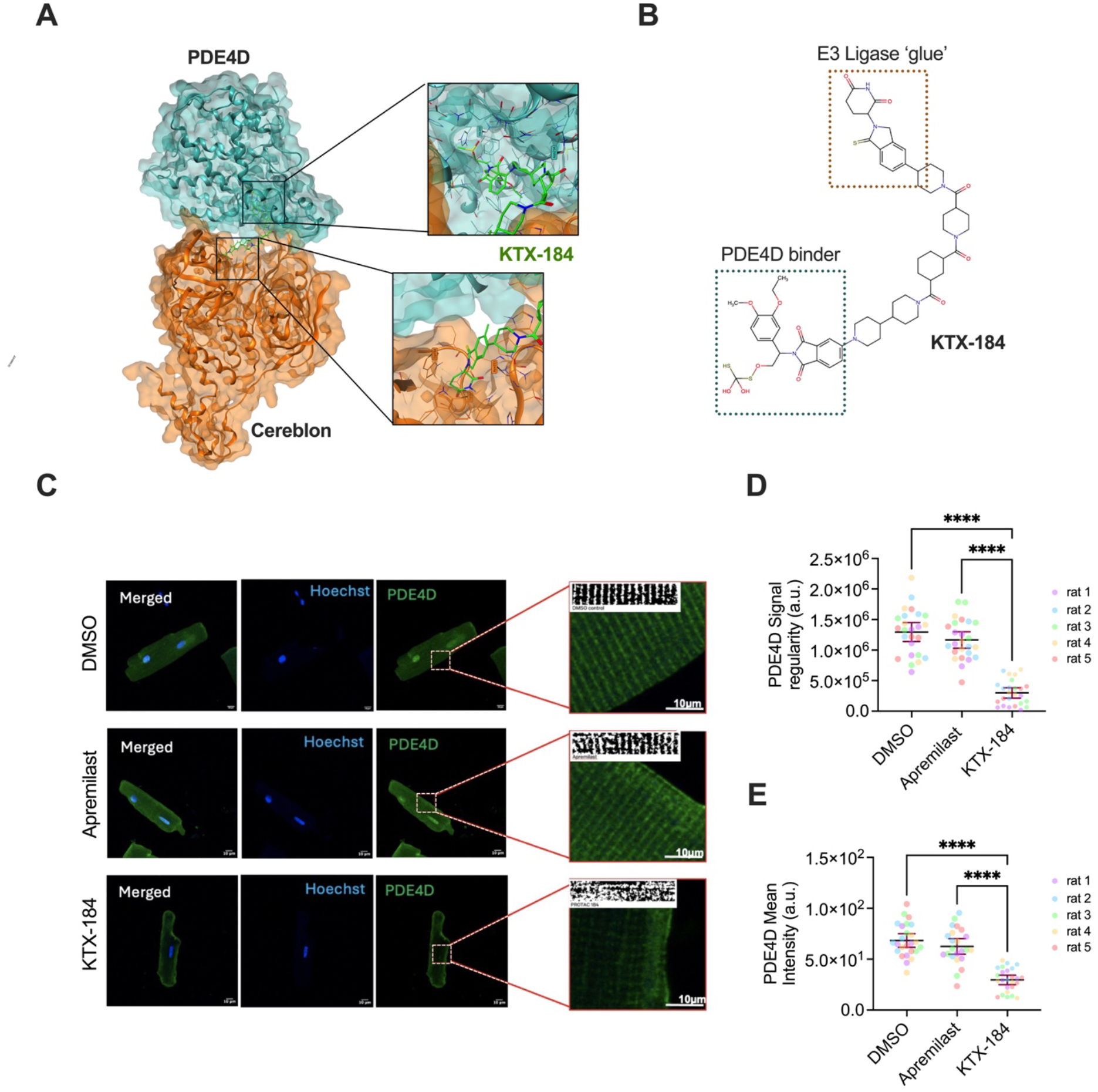
KTX-184 induces selective PDE4D degradation via cereblon-mediated targeted proteolysis. **a**, Three-dimensional molecular model of KTX-184 (green) docked into the tertiary complex with PDE4D (turquoise; PBD: 7CBQ) and cereblon (orange; PDB: 4CI1). Inset panels show site-view representations of computational docking. **b**, Two-dimensional skeletal structure of the KTX-184 with annotated functional domains: the PDE4D-binding warhead (blue dashed box) derived from the apremilast scaffold, and the cereblon-recruiting E3 ligase ‘glue’ moiety (orange dashed box), connected via a chemical linker. **c**, Representative confocal immunofluorescence images of adult rat left ventricular (LV) cardiomyocytes stained for PDE4D (green) and nuclear (Hoechst; blue) following 24-h treatment with vehicle (0.1% DMSO), apremilast (1 nmol/L), or KTX-184 (1 nmol/L). White dashed boxes indicate magnified regions shown at right; upper insets show binarised images used for Fourier transform-based regularity analysis. Scale bar, 10 µm. **d**, Quantification of PDE4D immunostaining signal regularity and **e**, mean fluorescence intensity in vehicle-(white bars, black points), apremilast-(light grey bars, blue points), and KTX-184-treated (dark grey bars, red points) cardiomyocytes. Individual data points are colour-coded by animal. Data are presented as mean ± S.E.M.; *n*= 23-25 cells from 5 animals. Statistical comparisons were made using a nested one-way ANOVA with Tukey’s multiple comparisons test. \*\*\*\**P*< 0.0001.

To assess the selectivity of KTX-184-mediated PDE4D degradation, we first characterised its activity across a range of cell systems. In H9c2 cardiomyoblasts, KTX-184 reduced PDE4D long-form protein abundance in a concentration-dependent manner (1 nmol/L-10 µmol/L), whilst PDE4B long-form expression was unaffected across the same concentration range (Supplementary Figure 1a-d). Immunofluorescence confirmed a significant reduction in cellular PDE4D abundance in KTX-184-treated H9c2 cells compared with apremilast-treated cells (Supplementary Figure 2a, b). Time-course analysis demonstrated that PDE4D long-form depletion was detectable as early as 2-h following KTX-184 treatment and was sustained through 24-h (Supplementary Figure 3a, b). Consistent with proteasome-mediated degradation, co-treatment with the proteasome inhibitor bortezomib fully prevented KTX-184-induced PDE4D depletion and was accompanied by accumulation of ubiquitinated proteins (Supplementary Figure 4a-c).

In adult rat LV cardiomyocytes, treatment with 1 nmol/L KTX-184 for 24-h disrupted the characteristic striated PDE4D immunostaining pattern observed in vehicle- and apremilast-treated cells (Figure 1c). Quantification of fluorescence regularity by Fourier transform-based analysis ^50^ and mean fluorescence intensity confirmed a significant reduction in PDE4D signal in KTX-184-treated cells, with no effect of apremilast at the same concentration (Figure 1d, e). Importantly, PDE4B immunostaining intensity and regularity were unaffected by KTX-184 (Supplementary Figure 5a-c), confirming isoform selectivity at nanomolar doses. Western blot analysis of rat LV cardiomyocyte lysates further demonstrated that KTX-184 significantly reduced total PDE4D and PDE4D5 protein abundance relative to apremilast (Supplementary Figure 6a-d).

### KTX-184 enhances β-adrenoceptor-induced cytosolic cAMP accumulation and PKA activity in rat left ventricular cardiomyocytes

FRET microscopy was used to examine the downstream effects of KTX-184 on β-AR signalling using two complementary biosensors: Epac1-camps, reporting cytosolic cAMP, and AKAR3-NES, reporting cytosolic PKA activity. Epac1-camps exhibited the expected diffuse cytosolic localisation in adult LV cardiomyocytes (Figure 2a, b). After 24-h pre-treatment with 1 nmol/L KTX-184, the magnitude of the isoprenaline-induced FRET response was markedly increased compared with vehicle- and apremilast-treated controls (Figure 2c), and quantification revealed a 2.8-fold increase in the β-AR-evoked cAMP response (Figure 2d). The effect of KTX-184 on cytosolic cAMP was robust across all six concentrations tested (10 pmol/L-1 µmol/L; Supplementary Figures 7 and 8), whereas apremilast augmented the FRET response only at 1 µmol/L (Supplementary Figure 8), consistent with its requirement for catalytic active-site occupancy to suppress enzymatic activity^51^ rather than targeted protein degradation^52^. To confirm that the enhanced cAMP response was dependent on proteasomal degradation of PDE4D, cardiomyocytes were co-treated with the proteasome inhibitor bortezomib (100 nmol/L). Bortezomib fully abolished the KTX-184-mediated enhancement of the isoprenaline-induced cAMP response, while having no significant effect on its own (Supplementary Figure 9), confirming that the functional effect of KTX-184 mechanistically depends on proteasomal degradation of PDE4D.

**Figure 2.**
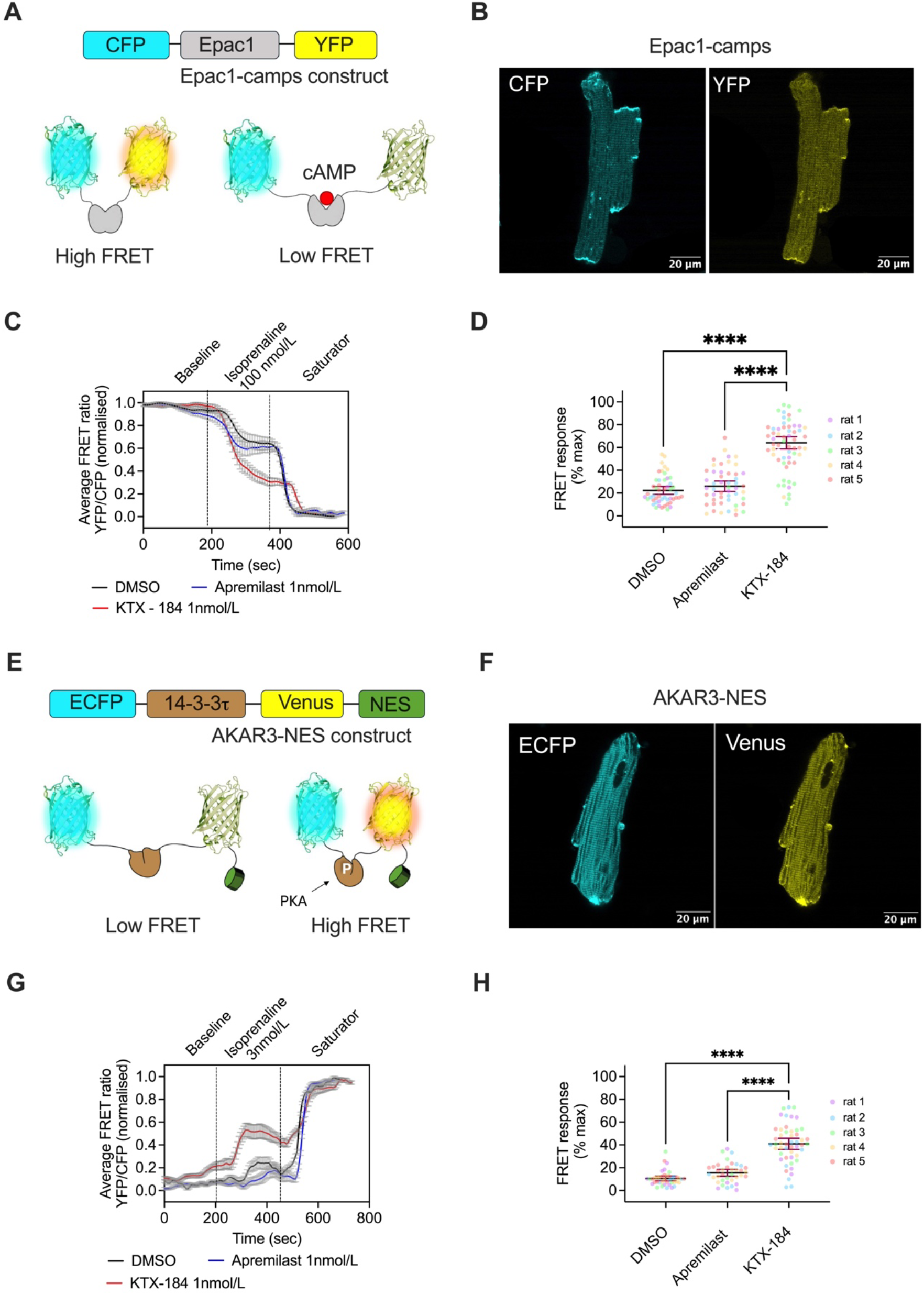
KTX-184 enhances cytosolic cAMP accumulation and PKA activity in adult rat left ventricular cardiomyocytes. **a**, Schematic of the Epac1-camps FRET-based cAMP biosensor. The construct comprises CFP and YFP fluorophores flanking the Epac1 cAMP-binding domain. cAMP binding induces a conformational change that reduces FRET efficiency (high FRET: unbound state; low FRET: cAMP-bound state). **b**, Representative pseudo-colour confocal images of adult rat LV cardiomyocytes expressing Epac1-camps, showing CFP (cyan) and YFP (yellow) fluorescence channels. Scale bars, 20 µm. **c**, Representative normalised Epac1-camps FRET ratio traces (YFP/CFP) from LV cardiomyocytes pre-treated for 24-h with vehicle (0.1% DMSO; grey), apremilast (1 nmol/L; blue), or KTX-184 (1 nmol/L; red). Cells were sequentially stimulated with isoprenaline (100 nmol/L) followed by a saturating cocktail of IBMX (100 µmol/L) and forskolin (10 µmol/L). Shaded areas represent S.D. **d**, Quantification of peak Epac1-camps FRET response expressed as a percentage of the maximal (saturator) response. Data are presented as mean ± 95% CI; *n*= 37-69 cells from 5 independent animals. Nested one-way ANOVA, with Tukey’s multiple comparisons test; \*\*\*\**P*< 0.0001; **e**, Schematic of the AKAR3-NES FRET-based PKA activity biosensor. The construct comprises ECFP and Venus fluorophores flanking a 14-3-3*τ* substrate sequence and nuclear export signal (NES). PKA-dependent phosphorylation of the substrate induces intramolecular binding to the 14-3-3*τ* domain, increasing FRET efficiency (low FRET: unphosphorylated; high FRET: PKA-phosphorylated). **f**, Representative pseudo-colour confocal images of adult rat LV cardiomyocytes expressing AKAR3-NES, showing ECFP (cyan) and Venus (yellow) fluorescence channels confirming cytosolic localisation. Scale bars, 20 µm. **g**, Representative normalised AKAR3-NES FRET ratio traces from LV cardiomyocytes pre-treated for 24-h with vehicle (0.1% DMSO; grey), apremilast (1 nmol/L; blue), or KTX-184 (1 nmol/L; red), stimulated with isoprenaline (3 nmol/L) followed by saturating IBMX (100 µmol/L) and forskolin (10 µmol/L). Shaded areas represent S.D. **h**, Quantification of peak AKAR3-NES FRET response expressed as a percentage of the maximal response. Data are presented as mean ± 95% CI; *n*= 37-49 cells from 5 independent animals. Nested one-way ANOVA; Tukey’s multiple comparisons test: \*\*\*\**P*< 0.0001.

To determine whether KTX-184-mediated PDE4D degradation translated increased cytosolic cAMP levels into enhanced downstream PKA signalling, cytosolic PKA activity was assessed using the AKAR3-NES biosensor, which localised to the cytosol as expected (Figure 2e, f). KTX-184 (1 nmol/L) significantly increased the isoprenaline-induced AKAR3-NES FRET response compared with both vehicle and apremilast controls (Figure 2g), with quantification demonstrating a 3.8-fold increase in cytosolic PKA activity relative to vehicle (Figure 2h). Apremilast at the same concentration had no significant effect on PKA activity, further supporting the functional superiority of targeted PDE4D degradation over catalytic inhibition at nanomolar concentrations^52,53^.

### KTX-184 does not modulate cardiomyocyte contractility, Ca^2^⁺ handling, or pro-contractile cAMP nanodomains

To determine whether KTX-184-mediated long-PDE4D degradation alters β-AR contractile responses, sarcomere shortening, Ca^2^⁺ transients, and contraction kinetics were examined in adult rat LV cardiomyocytes following 24-h pre-treatment with KTX-184 or apremilast at nanomolar and micromolar concentrations. Stimulation with isoprenaline produced the expected significant increases in sarcomere shortening and Ca^2^⁺ transient amplitude across all treatment groups (Figure 3a-d; Supplementary Figures 10, 11), consistent with positive inotropic and lusitropic effects^30,54^. Importantly, neither KTX-184 nor apremilast altered sarcomere shortening at either concentration tested (Figure 3a-d; Supplementary Figure 10), and no difference in Ca^2^⁺ transient amplitude or kinetics, including time to 90% peak contraction (TTP90) and time to 90% return to baseline (TTB90), was observed at micromolar or nanomolar concentrations (Supplementary Figure 11a-h). To assess potential pro-arrhythmic liability, the proportion of spontaneous extra beats was quantified in 1 Hz-paced cardiomyocytes over a 10 sec recording window. No significant difference was detected between KTX-184-, apremilast- and vehicle-treated cells at either concentration (Supplementary Figure 12a, b). Clinical studies have reported no QTc prolongation with the PDE4 inhibitor apremilast at therapeutic and supratherapeutic doses^55^, whereas the PDE3 inhibitor milrinone increased atrial arrhythmias in patients with decompensated heart failure^56^.

**Figure 3.**
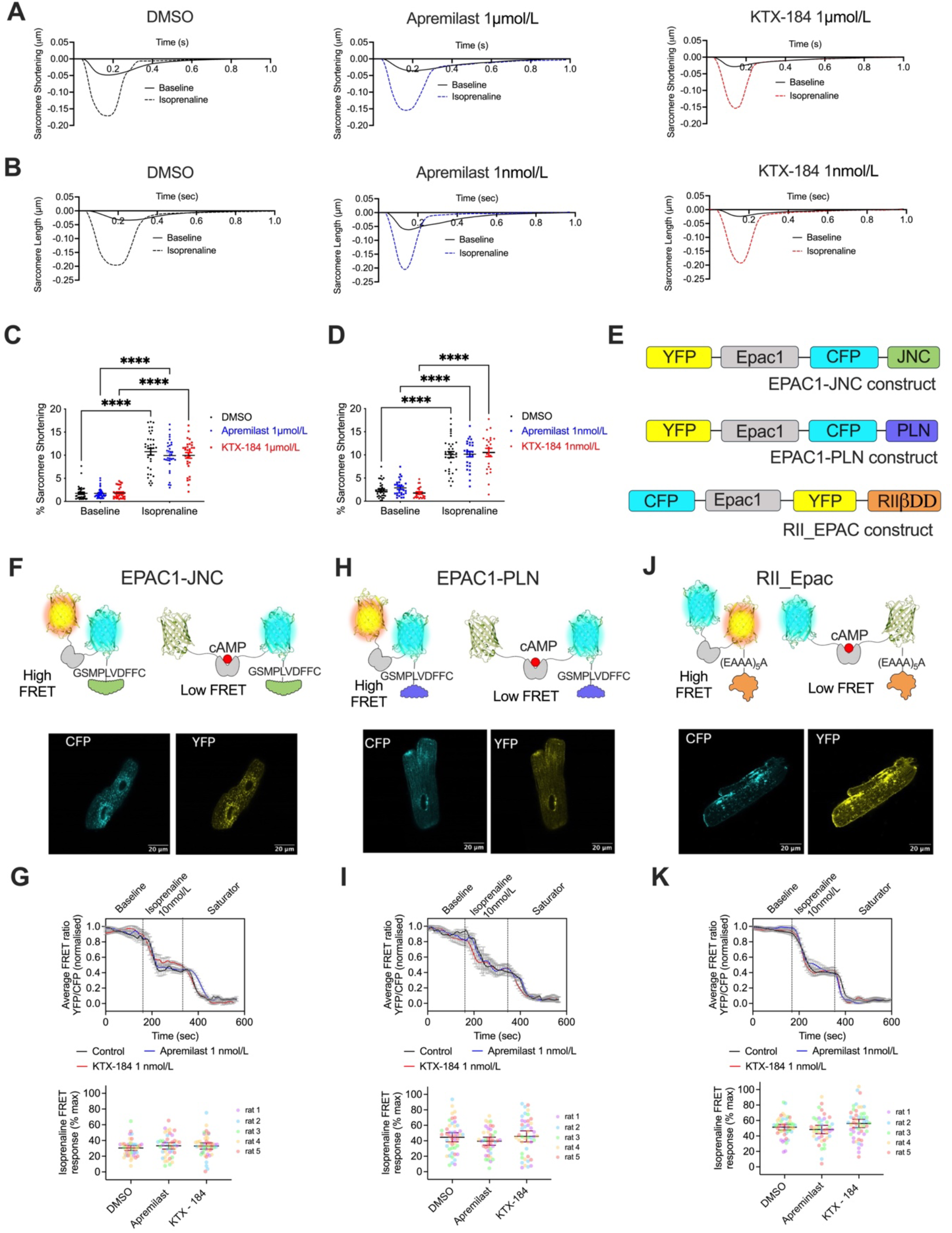
KTX-184 does not alter cardiomyocyte contractility or cAMP signalling within junctophilin, phospholamban-, or PKA-RII-associated signalling compartments. **a**, **b**, Representative sarcomere-shortening traces from adult rat left ventricular (LV) cardiomyocytes at baseline and following isoprenaline stimulation after 24-h pre-treatment with vehicle (0.1% DMSO), apremilast, or KTX-184 at micromolar (**a**) or nanomolar (**b**) concentrations. **c**, **d**, Quantification of percentage sarcomere shortening at baseline and following isoprenaline stimulation in cardiomyocytes pre-treated with micromolar (**c**) or nanomolar (**d**) concentrations of the indicated compounds. Data are presented as mean ± 95% CI; *n*= 33-45 cells from 5 animals. Two-way ANOVA with Tukey’s multiple comparisons test; baseline vs isoprenaline \*\*\*\**P*< 0.0001 for all groups. **e**, Schematics of the EPAC1-JNC, EPAC1-PLN, and RII_EPAC FRET-based cAMP biosensors, reporting cAMP dynamics within junctophilin (JNC)-, phospholamban (PLN)-, and PKA regulatory type II subunit (RII)-associated signalling compartments, respectively. **f**, **h**, **j**, Schematics illustrating the cAMP-dependent conformational change and FRET response of EPAC1-JNC (**f**), EPAC1-PLN (**h**), and RII_EPAC (**j**), with representative pseudo-colour confocal images of adult rat LV cardiomyocytes expressing each biosensor (CFP and YFP channels shown). Scale bars, 20 µm. **g**, **i**, **k**, Normalised FRET ratio traces (top; shaded areas represent S.D.) from cardiomyocytes pre-treated for 24-h with vehicle (0.1% DMSO; grey), apremilast (1 nmol/L; blue), or KTX-184 (1 nmol/L; red), stimulated with isoprenaline (10 nmol/L) followed by saturating IBMX (100 µmol/L) and forskolin (10 µmol/L), and quantification of the normalised isoprenaline-induced FRET response expressed as a percentage of the maximal response (bottom). Data are presented as mean ± 95% CI; *n* = 41-52 cells from 5 independent animals; nested one-way ANOVA with Tukey’s multiple comparisons test.

To determine whether the absence of contractile effects reflected preservation of pro-contractile cAMP nanodomains, localised FRET biosensors were used to probe cAMP dynamics within junctophilin-, phospholamban-, and PKA-RII-associated compartments. FRET responses from EPAC1-JNC (Figure 3f, g), EPAC1-PLN (Figure 3h, i), and RII_EPAC (Figure 3j, k) were indistinguishable between KTX-184-, apremilast-, and vehicle-treated cardiomyocytes, confirming that KTX-184 does not perturb cAMP signalling within contractile nanodomains. Consistent with these functional observations, western blot analysis revealed no change in phosphorylation of troponin I or phospholamban following KTX-184 treatment (Supplementary Figure 13a, b), confirming that PKA-mediated phosphorylation of key contractile targets is unaffected by long-PDE4D degradation.

### KTX-184 enhances plasma membrane-associated cAMP signalling via both β₁- and β₂-adrenoceptor subtypes

We next examined PDE4D-mediated regulation of plasma membrane-localised cAMP pools associated with distinct β-AR subtypes and sarcolemmal microdomains using a pmEpac2 FRET-based reporter ^57^. Previous studies have demonstrated that PDE4D regulates localised cAMP signalling at the plasma membrane by directly associating with β₁-AR prior to agonist stimulation and dissociating upon receptor activation, and by being recruited to β₂-AR via an agonist-dependent β-arrestin complex, thereby shaping spatially distinct β₁- and β₂-AR-mediated cAMP pools^58–63^. Here, we focused on β_1_- and β_2_-AR signalling, the major receptor subtypes controlling cardiomyocyte physiology ^64,65^ through spatially restricted cAMP pools ^2,38^ and differentially regulated by PDE4D ^58^. Adult LV cardiomyocytes expressing pmEpac2 displayed the expected plasma membrane localisation of CFP and YFP signal (Figure4a, b).

Following 24-h pre-treatment with 1 nmol/L KTX-184 or apremilast, only KTX-184 produced a significant increase in the isoprenaline-induced pmEpac2 FRET response, representing a 1.8-fold enhancement relative to vehicle control (Figure 4c). To dissect the contributions of individual β-AR subtypes, selective receptor antagonists were applied throughout stimulation. KTX-184 significantly enhanced cAMP responses mediated by both β₁-AR, assessed in the presence of the β₂-AR antagonist ICI 118,551 (Figure 4d), and β₂-AR, assessed in the presence of the β₁-AR antagonist CGP 20712A (Figure 4e). These findings were further corroborated using the cytosolic Epac1-camps biosensor under subtype-selective conditions, confirming that KTX-184-mediated enhancement of β₁- and β₂-AR-associated cAMP pools extends beyond the plasma membrane compartment (Supplementary Figure 14a, b).

**Figure 4.**
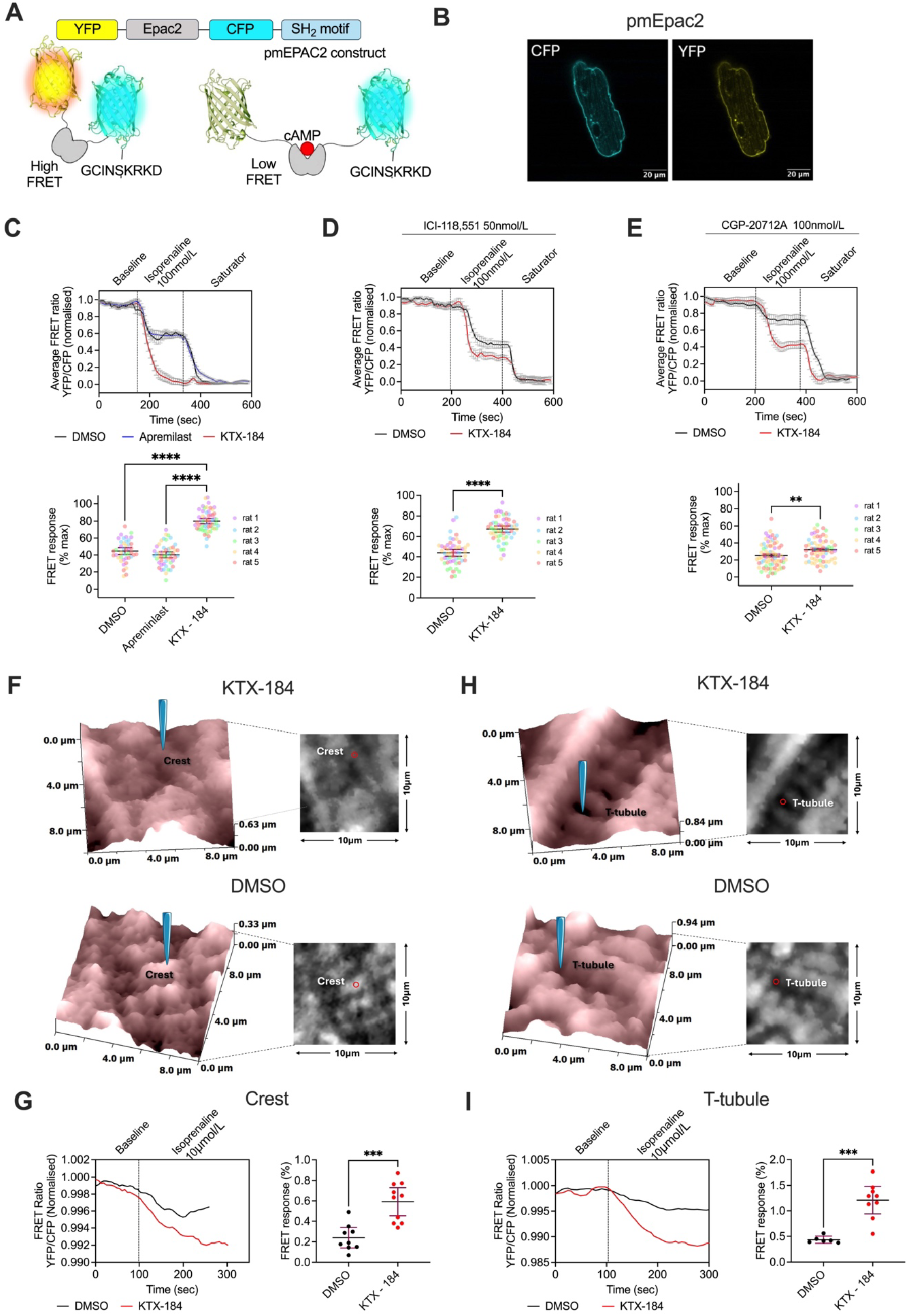
KTX-184 enhances plasma membrane-associated cAMP signalling in adult rat left ventricular cardiomyocytes. **a**, Schematic representation and operating principle of the plasma membrane-targeted pmEpac2 FRET-based cAMP biosensor. The construct is anchored to the plasma membrane via a SH_2_ motif. cAMP binding induces a conformational change resulting in decreased FRET efficacy. **b**, Representative pseudo-colour confocal images of an adult rat left ventricular (LV) cardiomyocytes expressing pmEpac2, showing CFP and YFP fluorescence channels. Scale bars, 20 μm. **c**, Normalised pmEpac2 FRET ratio traces (top), and quantification of the FRET response expressed as a percentage of the maximum response (bottom) in LV cardiomyocytes following 24-h pre-treatment with vehicle (0.1% DMSO; grey), apremilast (1 nmol/L; blue), or KTX-184 (1 nmol/L; red). Cells were sequentially stimulated with isoprenaline (100 nmol/L) and saturating 3-isobutyl-1-methylxanthine (IBMX; 100 μmol/L) plus forskolin (10 μmol/L). Data presented as mean ± 95% CI, *n* = 43 - 56, from 5 rats. Nested one-way ANOVA; Tukey’s multiple comparison test: \*\*\*\**P*<0.0001. **d**, **e**, Normalised pmEpac2 FRET ratio traces (top) and quantification of FRET responses (bottom) following 24-h pre-treatment with vehicle (0.1% DMSO) or KTX-184 (1 nmol/L). Cells were stimulated with isoprenaline in the presence of ICI-118,551 (50 nmol/L; **d**) or CGP-20712A (100 nmol/L; **e**), followed by saturating agonists. Nested unpaired t-test, \*\**P*=0.0052. **f**, Representative scanning ion conductance microscopy (SICM) topography maps of LV cardiomyocytes treated with KTX-184 (1 nmol/L) or vehicle (0.1% DMSO), illustrating positioning of the SICM nanopipette over a sarcolemmal crest region for localised isoprenaline (10 μmol/L) application. **g**, Representative pmEpac2 FRET ratio traces during localised sarcolemmal crest stimulation (left) and quantification of the locally evoked FRET response (right) in vehicle- and KTX-184-treated cardiomyocytes. Data are mean ± 95% CI, *n* = 9 - 10, from 5 rats Welch’s t-test: \*\*\**P*=0.0002. **h**, Representative SICM topography maps illustrating nanopipette positioning over a T-tubule opening for localised isoprenaline application in KTX-184- and vehicle-treated cardiomyocytes. **I**, Representative pmEpac2 FRET ratio traces during localised T-tubule stimulation (left) and quantification of the locally evoked FRET response (right). Data are mean ± 95% CI, *n* = 6 - 9, from 5 rats; Welch’s t-test: \*\*\**P*=0.0001.

To determine whether the spatial localisation of β-AR stimulation influenced the magnitude of the KTX-184 effect, localised isoprenaline application was performed via SICM nanopipettes to either the sarcolemmal crest or T-tubule openings of pmEpac2-expressing cardiomyocytes. Following localised stimulation at sarcolemmal crests, KTX-184-treated cardiomyocytes exhibited a 2.2-fold greater FRET response than vehicle-treated controls (Figure 4f, g). Similarly, localised stimulation at T-tubule openings produced a significantly enhanced cAMP response in KTX-184-treated cells relative to vehicle (Figure 4h, i), demonstrating that KTX-184 augments β-AR-evoked cAMP signalling at both major sarcolemmal microdomains.

### KTX-184 attenuates AngII-induced hypertrophic remodelling and enhances T-tubule-associated nuclear PKA signalling in rat left ventricular cardiomyocytes

To assess the functional consequences of the localised KTX-184 effect on plasma membrane-localised cAMP pools, we tested whether targeted PDE4D degradation could prevent AngII-induced hypertrophic remodelling, given prior evidence of anti-hypertrophic effects following PDE4D ablation^20,21^. KTX-184 preserved cardiomyocyte morphology under hypertrophic conditions (Figure 5a), reversing the AngII-induced increase in cell width and reduction in aspect ratio (Figure 5b), without significantly affecting cell length (Supplementary Figure 15). A comparable anti-hypertrophic effect was observed in neonatal rat ventricular cardiomyocytes, where KTX-184 significantly reduced AngII-induced increases in cell area, whilst apremilast at the same concentration had no effect (Supplementary Figure 16).

**Figure 5.**
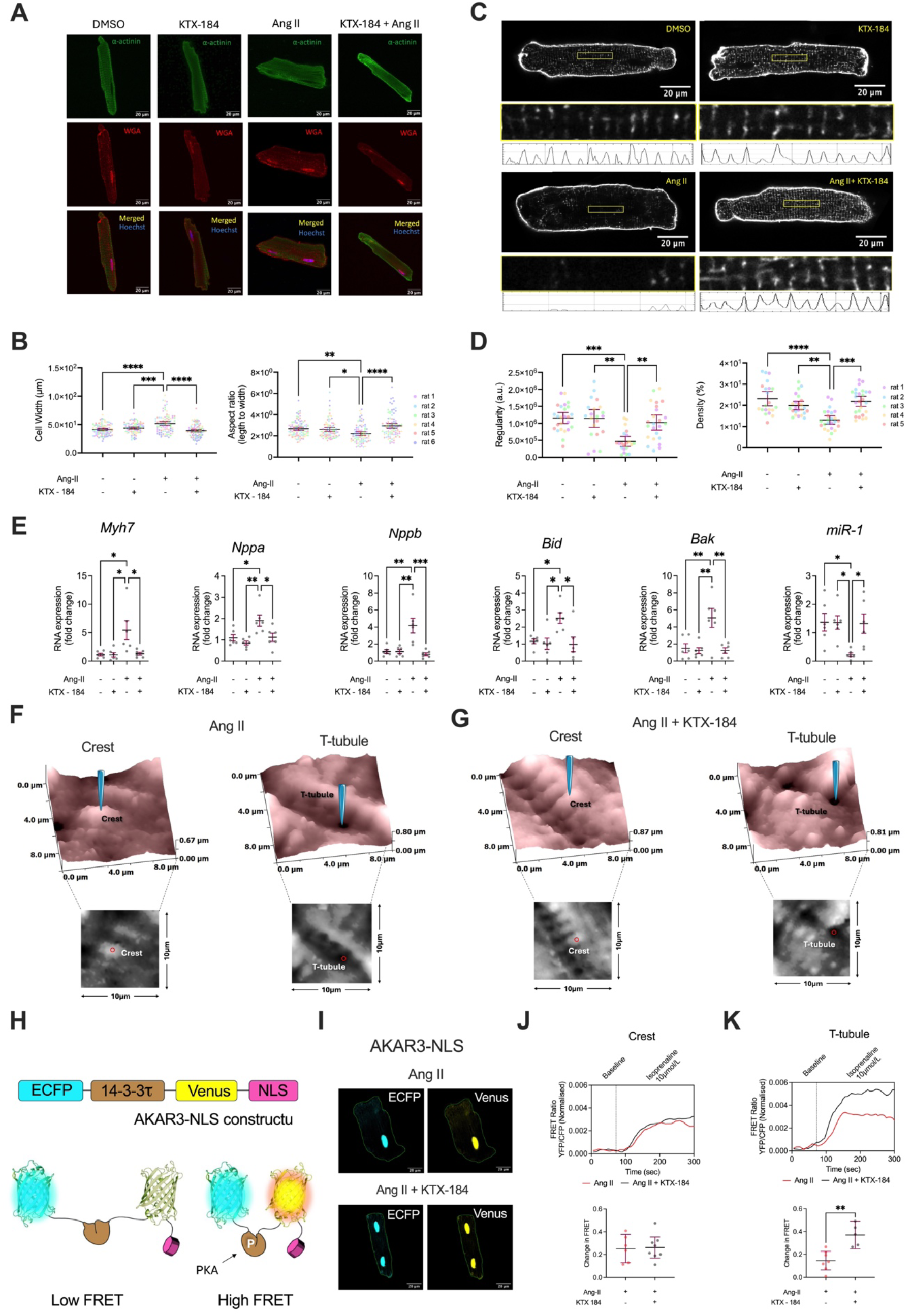
KTX-184 attenuates AngII-induced cardiomyocyte hypertrophy and enhances T-tubule-associated cAMP/PKA signalling to the nuclear compartment. **a**, Representative pseudo-colour confocal images of adult rat left ventricular (LV) cardiomyocytes treated with vehicle (0.1% DMSO), KTX-184 (1nmol/L), angiotensin II (AngII; 20 μmol/L), or AngII with KTX-184, labelled for α-actinin (green) and wheat germ agglutinin (WGA; red), nuclei counterstained with Hoechst. Scale bars, 20 μm. **b**, Quantification of cardiomyocyte width (left) and aspect ratio (right) under the indicated treatment conditions. Data presented as mean ± 95% CI, *n* = 84 - 97, from 6 rats. Nested one-way ANOVA with Tukey’s multiple comparison test: \**P <*0.05, \*\**P <* 0.01, \*\*\**P <* 0.001, \*\*\*\**P <* 0.0001. **c**, Representative Di-8-ANEPPS-labelled images of adult rat LV cardiomyocytes under the indicated treatment conditions, with enlarged regions T-tubule organisation and corresponding fluorescence intensity profiles. Scale bars, 20 μm. **d**, Quantification of T-tubule regularity (left) and density (right) under the indicated treatment conditions. Data are mean ± 95% CI, *n* = 21-25, from 5 rats. Nested one-way ANOVA with Tukey’s multiple comparison test: \*\**P <* 0.01, \*\*\**P <* 0.001, \*\*\*\**P <* 0.0001. **e**, Relative mRNA expression of the hypertrophy-associated genes (*Myh7*, *Nppa*, *Nppb*), apoptosis-associated genes (*Bid, Bak*), and *miR-1* following treatment with AngII in the presence or absence of KTX-184, expressed as fold change relative vehicle control. Data presented as mean ± S.E.M., *n* = 6 rats. \**P* ≤ 0.05, \*\**P* ≤ 0.01, \*\*\**P* ≤ 0.001 based on the one-way ANOVA with Tukey’s multiple comparison test.. **f**, **g**, Representative SICM topography maps of adult rat LV cardiomyocytes treated with AngII (**f**) or AngII in combination with KTX-184 (**g**), illustrating nanopipette positioning over a sarcolemmal crest (left) or T-tubule opening (right) for local β-adrenergic stimulation. **h**, Schematic representation and operating principle of the nuclear-targeted AKAR3-NLS FRET-based PKA activity biosensor. PKA-dependent phosphorylation of the substrate induces a conformational change resulting in increased FRET. **i**, Representative pseudo-colour confocal images of adult rat LV cardiomyocytes expressing AKAR3-NLS following treatment with AngII or AngII combined with KTX-184, showing ECFP and Venus fluorescence. Scale bars, 20 μm. **j**, **k**, Representative AKAR3-NLS FRET ratio traces (top) and quantification of nuclear PKA responses (bottom) following local β-adrenergic stimulation at a sarcolemmal crest (**j**) or T-tubule opening (**k**) in cardiomyocytes under indicated treatment conditions. Data are mean ± 95% CI. **j**: *n* = 6-8, 4-5 rats, unpaired Welch’s *t*-test; **k**: *n* = 5-7 from 4-5 rats, ** *P*=0.0032.

KTX-184 also preserved T-tubule organisation under hypertrophic conditions, reversing the AngII-mediated reduction in both T-tubule regularity and density (Figure 5c, d). Consistent with attenuation of the hypertrophic programme ^66–70^, KTX-184 reversed the AngII-induced upregulation of the hypertrophy-associated genes *Myh7*, *Nppa*, and *Nppb*, the pro-apoptotic genes *Bid* and *Bak*, and restored *miR-1* expression towards control levels (Figure 5e).

Given that β-AR stimulation and cAMP-dependent nuclear PKA activity drive hypertrophic gene transcription^7,71^, we next assessed whether KTX-184 modulates nuclear PKA signalling using the AKAR3-NLS biosensor (Figure 5f). Under non-hypertrophic conditions, KTX-184, but not apremilast, significantly enhanced nuclear PKA activity in response to isoprenaline stimulation (Supplementary Figure 17). In AngII-treated cardiomyocytes, whole-cell FRET recordings demonstrated that KTX-184 selectively increased nuclear PKA activity and plasma membrane-associated cAMP responses, whilst leaving global cytosolic cAMP responses unchanged (Supplementary Figure 18a-c). To probe spatially resolved signalling, localised β-adrenergic stimulation was applied via SICM nanopipettes to either sarcolemmal crests or T-tubule openings in cardiomyocytes treated with AngII alone or a combination of AngII and KTX-184 (Figure 5g). KTX-184 selectively enhanced the T-tubule-associated nuclear PKA response in hypertrophic cardiomyocytes (Figure 5i), whilst the crest-associated nuclear response was unaffected (Figure 5h). This spatial selectivity was further confirmed by pmEpac2 measurements, which revealed that KTX-184 augmented membrane-associated cAMP responses specifically at T-tubule sites, but not at crest regions, in AngII-treated cells (Supplementary Figure 19). As the PDE4D-HSP20 interaction has been previously implicated in the regulation of cardioprotective PKA signalling^72,73^, we examined whether KTX-184-mediated PDE4D degradation altered HSP20 phosphorylation. No significant change in phospho-HSP20 levels was detected in adult rat LV cardiomyocytes following 24-h KTX-184 treatment relative to vehicle or apremilast controls (Supplementary Figure 20), suggesting that the anti-hypertrophic effect of KTX-184 is not mediated by acute changes in HSP20 phosphorylation under basal conditions.

### KTX-184 enhances β-adrenoceptor-induced cytosolic cAMP release in isolated human donor cardiomyocytes and attenuates hypertrophic remodelling in human myocardial tissue slices

To determine whether the effects of KTX-184 observed in rat LV cardiomyocytes translate to human LV myocardium, we evaluated 1 nmol/L KTX-184 in isolated human donor LV cardiomyocytes. In human LV myocardium, PDE4D has been shown to be expressed and enzymatically active^19^ and to be upregulated in hypertrophy and heart failure^20^. To establish the appropriate isoprenaline concentration for FRET microscopy on human donor cardiomyocytes, Epac1-camps-expressing cells (Figure 6a) were stimulated with 10 nmol/L and 100 nmol/L isoprenaline. The 100 nmol/L dose produced a response approaching sensor saturation, whereas 10 nmol/L elicited a significantly smaller response (Figure 6b), providing sufficient dynamic range to detect the modulatory effect of KTX-184 and apremilast.

**Figure 6.**
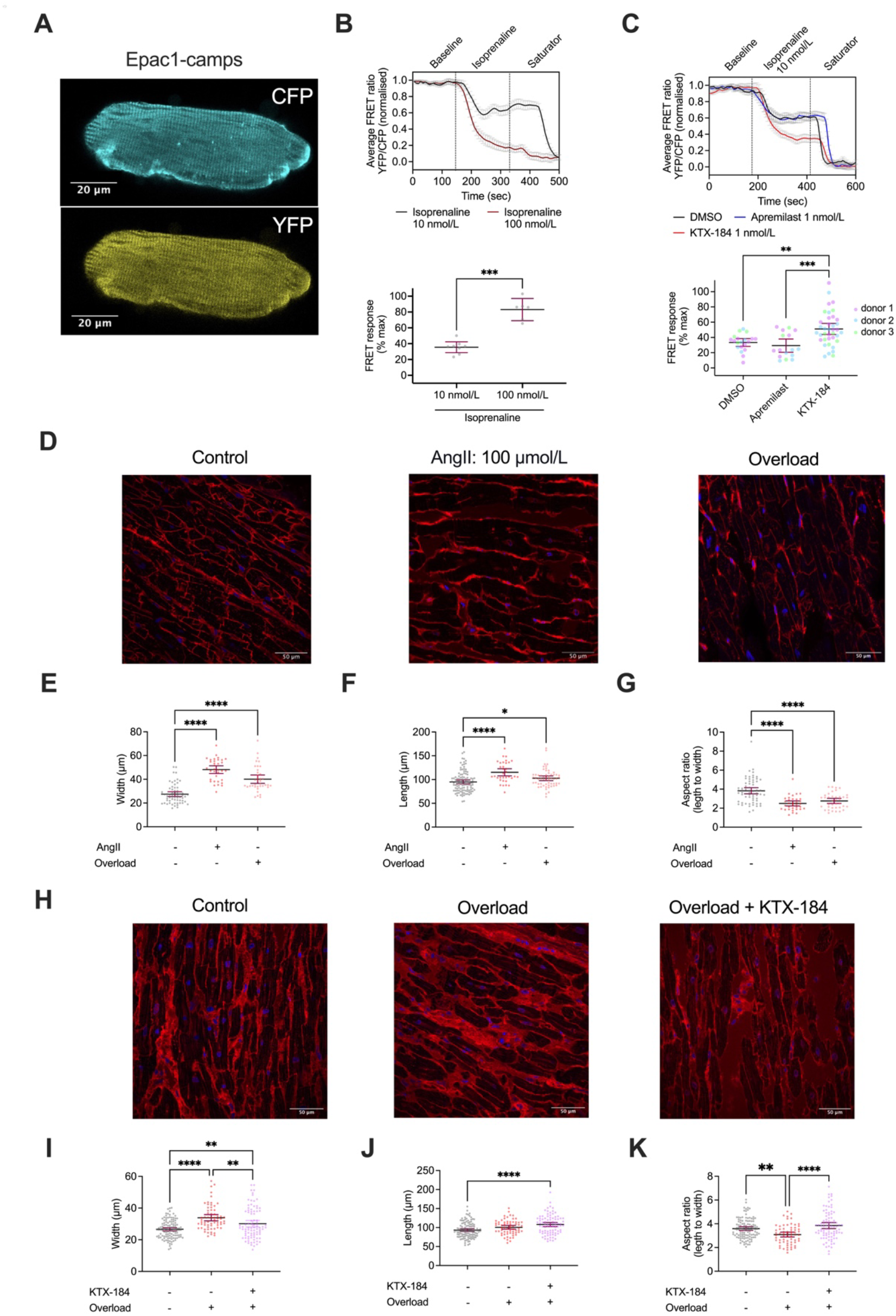
KTX-184 enhances global cAMP signalling in human left ventricular cardiomyocytes and attenuates hypertrophic remodelling in human myocardial tissue. **a**, Representative pseudo-colour confocal images of adult human donor left ventricular (LV) cardiomyocytes expressing the Epac1-camps FRET-based cAMP biosensor, showing CFP and YFP fluorescence. Scale bars, 20 μm. **b**, Normalised Epac1-camps FRET ratio traces (top) and quantification of the FRET responses a percentage of the maximum response (bottom) following stimulation with isoprenaline at 10 nmol/L or 100 nmol/L, followed by saturating agonist (the 3-isobutyl-1-methylxanthine [IBMX, 100 μmol/L], forskolin [10 μmol/L]) demonstrating concentration-dependent cAMP responses in human LV cardiomyocytes. Data presented as mean ± 95% CI, *n* = 6-8, Mann Whitney test, \*\*\**P*=0.0007 **c**, Normalised Epac1-camps FRET ratio traces (top) and quantification of the FRET response (bottom) in human LV cardiomyocytes following 24-h pre-treatment with vehicle (0.1% DMSO; grey), apremilast (1 nmol/L; blue) or KTX-184 (1 nmol/L; red), stimulated sequentially with isoprenaline (10 nmol/L) and saturating IBMX (100 μmol/L) plus forskolin (10 μmol/L). Data presented as mean ± 95% CI, and *n* = 16 - 28 from 3 donors. Kruskal-Wallis test, Dunn’s multiple comparison: \*\**P*=0.0038, \*\*\**P*=0.0010. **d**, Representative images of human donor myocardial slices maintained for 72-h under physiological preload (control), treated with angiotensin II (AngII; 100 μmol/L), or subjected to mechanical overload. Cardiomyocyte boundaries were visualised using wheat germ agglutinin (WGA; red), with nuclei counterstained with DAPI (blue). Scale bars, 50 μm. **e**-**g**, Quantification of cardiomyocyte width (**e**), length (**f**) and aspect ratio (**g**) under control, AngII-treated, and mechanical-overload conditions, demonstrating comparable morphological remodelling induced by both stimuli. Data are mean ± 95% CI; *n* = 34 - 117 from 3 donors. Welch ANOVA test with Dunnett’s multiple comparison, \**P* <0.05, \*\**P* <0.01, \*\*\**P* <0.001 and \*\*\*\**P* <0.0001 **h**, Representative images of human donor myocardial slices under physiological preload (control) or mechanical overload for 72 h in the absence or presence of KTX-184 (100 nmol/L). Scale bars, 50 μm. **i**-**k**, Quantification of cardiomyocyte width (**i**), length (**j**) and aspect ratio (**k**), demonstrating attenuation of overload-induced morphological remodelling by KTX-184. Data are presented as mean ± 95% CI, *n* = 65-118 cardiomyocytes from 3 donors, \**P* ≤ 0.05, \*\**P* ≤ 0.01, \*\*\**P* ≤ 0.001 and \*\*\*\**P* ≤ 0.0001, based on Welch ANOVA with Tukey’s multiple-comparisons test.

Following 24-h pre-treatment with 1 nmol/L KTX-184 or apremilast, the cytosolic cAMP response to 10 nmol/L isoprenaline was assessed using Epac1-camps FRET microscopy (Figure 6c). KTX-184 significantly enhanced the FRET response by 57% relative to vehicle, whilst apremilast had no significant effect at the same concentration (Figure 6c), consistent with the pattern observed in rat cardiomyocytes.

To assess the translational relevance of these findings in a more physiologically representative human cardiac model, we employed human donor myocardial tissue slices^47^. We first validated the hypertrophic phenotype by comparing morphological remodelling induced by mechanical overload and AngII stimulation (Figure 6d). Both conditions produced comparable increases in cardiomyocyte width (Figure 6e), length (Figure 6f), and altered aspect ratio (Figure 6g), confirming that AngII treatment under physiological preload recapitulates the overload-induced remodelling phenotype in this model. We then assessed the effect of KTX-184 in slices subjected to mechanical overload for 72 h (Figure 6h). KTX-184 treatment significantly reduced cardiomyocyte width and increased the aspect ratio in overloaded slices compared with untreated overloaded slices (Figure 6i, k), despite increased cell length in KTX-184-treated overloaded slices (Figure 6j), indicating at least partial attenuation of overload-induced hypertrophic remodelling in human myocardial tissue.

## DISCUSSION

To the best of our knowledge, this study describes the first example of targeted degradation of long PDE4D isoforms by the PROTAC KTX-184. We show that this approach can precisely reshape β-AR/cAMP signalling in adult ventricular cardiomyocytes to attenuate hypertrophic remodelling without affecting Ca^2+^ dynamics that regulate heartbeat. By eliminating a compartmentalised pool of PDE4D, KTX-184 amplifies β-adrenergic-evoked cAMP and PKA signalling in discrete subcellular nanodomains, most prominently at the plasma membrane and nuclear envelope, without disturbing excitation-contraction coupling or Ca^2^⁺ handling. This spatial precision distinguishes targeted PDE4D proteolysis from standard catalytic PDE4 inhibition and offers a strategy to promote cardioprotective signalling while avoiding inotropic or arrhythmogenic side effects.

Our findings align with and extend a rich body of work demonstrating that cAMP/PKA signalling in cardiomyocytes is highly compartmentalised. Using FRET reporters, we previously showed that β_1_-AR stimulation generates global cAMP elevations, whereas β_2_-AR signals are locally confined, and that this confinement reflects compartmentalised PDE activity and membrane architecture rather than Gᵢ coupling^2^. Subsequent biochemical work revealed that this receptor specificity is enforced by isoform-specific PDE4D interactions: PDE4D8 pre-associates with β_1_-AR and dissociates upon agonist stimulation, whereas PDE4D5 is recruited to β_2_-AR via an agonist-dependent β-arrestin complex ^58,62^. This pre-assembled PDE4D8 “gatekeeper” protects unoccupied β_1_-ARs from PKA-dependent desensitisation and, upon agonist binding, dissociates to permit a rapid local rise in cAMP ^58^. In parallel, Di Benedetto and colleagues demonstrated that PKA-I and PKA-II define distinct intracellular signalling compartments tethered by AKAPs, with PDE2 and PDE4 providing non-redundant control of cAMP hydrolysis ^6^. Consistent with this specialised regulation, our unchanged RII_Epac, Epac1-JNC and Epac1-PLN signals indicate preserved PKA-II/contractile microdomains, while increased AKAR-NLS and pmEpac2 responses reflect relief of nuclear and sarcolemmal constraints. Long PDE4D isoforms anchored to mAKAPβ ^63^at the nuclear envelope^74^ insulate β_2_-ARs from activating nuclear PKA and gene transcription, thereby enforcing receptor-specific transcriptional outcomes; β_1_-ARs overcome this barrier in part via catalytic-subunit diffusion to the nucleus ^7^. Together, these studies underscore how PDE4D isoforms, anchored to distinct receptors and scaffolds, serve as gatekeepers of receptor-to-nucleus second messenger signalling within cardiomyocytes.

Recent gene-therapy studies strengthen this interpretation by mapping PDE4 subfamily control across Ca^2^⁺-handling microdomains *in vivo* ^60^. In mice, targeted FRET probes revealed that PDE4D predominates at SERCA2a, whereas the RyR2 microdomain is PDE4B-centric, with both PDE4B and PDE4D shaping caveolin-rich plasma membrane cAMP dynamics ^60,61^. AAV9-PDE4B3 overexpression restored altered cAMP microdomains, reduced Ca^2^⁺ leak and arrhythmias, and attenuated pressure-overload remodelling, consistent with close PDE4B proximity to RyR2 ^39,40,60^. PDE4D anchoring to distinct sarcolemmal and nuclear scaffolds further shapes receptor-specific cAMP pools^59,75^, with PDE4D being specifically required for β₂-AR subtype-selective signalling in cardiomyocytes^61^. Changes in sarcolemmal and nuclear microdomains have been directly linked to disease progression in vivo^41^. In our study, KTX-184 selectively augmented pmEpac2 and AKAR-NLS signals but did not change Epac1-PLN or E1-JNC readouts. The absence of a RyR2-adjacent effect of PDE4D manipulation mediated by KTX-184 likely reflects the predominance of PDE4B in that compartment^39,60^, which is desirable as it avoids increasing arrhythmic risk^40^. Similarly, the unchanged Epac1-PLN signal following KTX-184 treatment suggests preferential depletion of long PDE4D isoforms at sarcolemmal and nuclear scaffolds, while preserving PLN/RyR2/PKA-RII nanodomains ^41,60^, consistent with the absence of inotropic or Ca^2^⁺-transient changes in our preparations.

The clinical use of PDE3 inhibitors such as milrinone to restore impaired β-adrenergic responsiveness in heart failure has been limited by increased ventricular arrhythmias and mortality^56,76^ in patients with HFrEF^17,10^ attributable to dysregulated cAMP signalling and abnormal sarcoplasmic reticulum Ca^2^⁺ handling^77^. In contrast, PDE4 inhibition appears mechanistically distinct, regulating highly compartmentalised cAMP nanodomains rather than directly enhancing global inotropy^18–21^. Importantly, KTX-184-mediated degradation of long PDE4D isoforms enhanced β-adrenergic/cAMP signalling without altering Ca^2^⁺ transients, sarcomere shortening, or arrhythmic events, suggesting that selective targeting of PDE4D may preserve cardioprotective signalling whilst avoiding the pro-arrhythmic liabilities associated with PDE3 inhibition.

KTX-184 bypasses PDE4D-mediated cAMP braking, allowing β-AR stimulation to engage transcriptional PKA pathways without altering Ca^2^⁺ transients or contractility. This mechanistic selectivity is critical, as excessive activation of contractile PKA targets, such as phospholamban or troponin I, can drive maladaptive remodelling^71^. By targeting nuclear and perinuclear long PDE4D isoforms^7,75^, KTX-184 may instead promote phosphorylation of substrates with anti-hypertrophic and anti-apoptotic actions, though the precise downstream mechanisms warrant further investigation. This interpretation is supported by consistent findings across complementary *in vitro* rat and *ex vivo* human models, including human myocardial slices that preserve native multicellularity, extracellular matrix, tissue architecture, and biomechanical load^47^.

These findings align with emerging evidence that PDE4D contributes to pathological remodelling through multiple mechanisms extending beyond cAMP compartmentation. While β₁-AR-evoked nuclear PKA can raise inducible cAMP early repressor (ICER) ^7^, scaffold context and PDE4D gating determine transcriptional bias; selectively relieving long PDE4D isoforms favours CREB-SIRT1-linked cardioprotective programmes^20,21^. PDE4D upregulation suppresses CREB-SIRT1 signalling and PINK1/Parkin-mediated mitophagy in failing hearts, promoting mitochondrial dysfunction, reactive oxygen species accumulation, and hypertrophy, whilst selective PDE4D inhibition or haploinsufficiency restores mitophagy, though complete ablation is maladaptive owing to loss of RyR2-associated PDE4D3^21^. PDE4D upregulation also depresses cAMP-PKA-CREB-SIRT1 signalling in cardiomyocytes and fibroblasts, reducing SERCA2a expression and miR-1, a muscle-enriched microRNA restraining hypertrophic gene programmes including MEF2a and GATA4^20,78,79^, and driving hypertrophy and fibrosis via TGF-β1/Smad signalling; PDE4D inhibition or knockdown restored SERCA2a/miR-1 and reduced remodelling^20^, and has similarly been linked to improved SERCA2a-associated Ca^2^⁺ handling in diabetic cardiomyopathy^80^. Additionally, the perinuclear mAKAP-PKA-PDE4D3-Epac1 complex enables cAMP-dependent activation of Epac1-Rap1 to suppress pro-hypertrophic ERK5 signalling, suggesting a potential complementary mechanism for the anti-hypertrophic effects of KTX-184 alongside its enhancement of nuclear PKA activity^81,82^. Together, these data extend the PDE4D axis upstream of excitation-contraction coupling to integrate mitochondrial, sarcoplasmic, and fibrotic mechanisms. In this context, long-PDE4D degradation by KTX-184 could provide multiple benefits: freeing nuclear and membrane cAMP pools to normalise transcription, restoring CREB-SIRT1 programmes linked to mitophagy and SERCA2a/miR-1 homeostasis, consistent with the KTX-184-mediated reversal of AngII-induced miR-1 downregulation observed here, whilst sparing RyR2/PLN nanodomains and thus avoiding inotropy^20,21^. KTX-184-mediated augmentation of nuclear PKA activity may also promote HDAC5 phosphorylation and nuclear retention, thereby repressing MEF2-dependent hypertrophic transcription and potentially contributing to the suppression of fetal gene expression, including Nppa and Nppb ^83^.

The preservation of T-tubule organisation is likely to stabilise β_2_-AR/cAMP nanodomains that depend on intact T-tubule and caveolar architecture and are redistributed in heart failure ^3,4,38^. Localised SICM stimulation confirmed larger β-AR cAMP gains at both T-tubule and crest sites after KTX-184, consistent with nanodomain-specific relief of PDE4D braking and with prior demonstrations that modulation of cAMP nanodomains profoundly affects hypertrophic remodelling ^2,6,7^. KTX-184 achieves these protective effects without global PDE4 inhibition, which is associated with emesis, possible arrhythmias, and contractile dysregulation^75^. Recent PDE4 PROTACS developed using PDE4 inhibitor warheads have yielded varied selectivity profiles: compound “9m” shows moderate PDE4B selectivity but very low potency (DC_50_ ∼42 µmol/L), while KTX-207 and BTX-AP04 are highly potent and selective for short-form PDE4 isoforms^23,48,84^, which are upregulated during inflammation to promote T-cell activation and represent attractive targets for autoimmune diseases such as atopic dermatitis. Here, we report the first PDE4 long-form-selective degrader with potent anti-hypertrophic activity in the heart, illustrating the unparalleled isoform selectivity achievable through PROTAC engineering ^85^.

Limitations of this study include its reliance on isolated cardiomyocyte systems; *in vivo* efficacy, pharmacokinetics, and safety of KTX-184 remain to be established. Whilst our data indicate selective degradation of long PDE4D isoforms, isoform-resolved proteomics and interactome mapping will be needed to confirm which complexes are disrupted. Transcriptomic profiling may further define the full spectrum of gene programs rebalanced by this intervention.

In conclusion, KTX-184 exemplifies how selective, targeted protein degradation of PDE4 can rewire cAMP nanodomains in the heart. By eliminating long PDE4D isoforms tethered at the plasma membrane and nuclear scaffolds, KTX-184 enhances receptor-to-nucleus β-AR signalling and may liberate cardioprotective PKA phosphorylation, thereby attenuating hypertrophic remodelling without disturbing contractility. These findings suggest that PROTAC-mediated PDE4D degradation may provide a new avenue for precision therapy in cardiac hypertrophy and heart failure, distinct from catalytic inhibition strategies.

## CLINICAL RELEVANCE

Hypertrophic cardiomyopathy is a condition in which areas of the heart muscle become stiff and thickened, progressively impairing cardiac function. Current therapies largely focus on symptom management and reducing hemodynamic burden rather than directly targeting the molecular mechanisms that drive pathological remodelling. Here, we provide the first evidence that compartmentalised regulation of cyclic-AMP signalling through targeted degradation of long PDE4D isoforms can attenuate maladaptive hypertrophic gene activation and structural remodelling of cardiac tissue. Importantly, unlike PDE3 inhibitors, whose clinical use in heart failure has been limited by pro-arrhythmic effects linked to dysregulated Ca^2^⁺ handling, KTX-184 enhanced β-adrenergic/cAMP signalling without altering Ca^2^⁺ transients or increasing arrhythmic activity, suggesting a potentially safer strategy for targeting maladaptive cardiac signalling.

## ACKNOWLEDGMENTS

The authors thank the Center of Excellence Cellular Mechanosensing and Functional Microscopy at Imperial College London.

