## Supplemental data for "Long-PDE4D degradation attenuates left ventricular cardiomyocyte hypertrophic remodelling"

** - corresponding authors*

**Corresponding authors:**

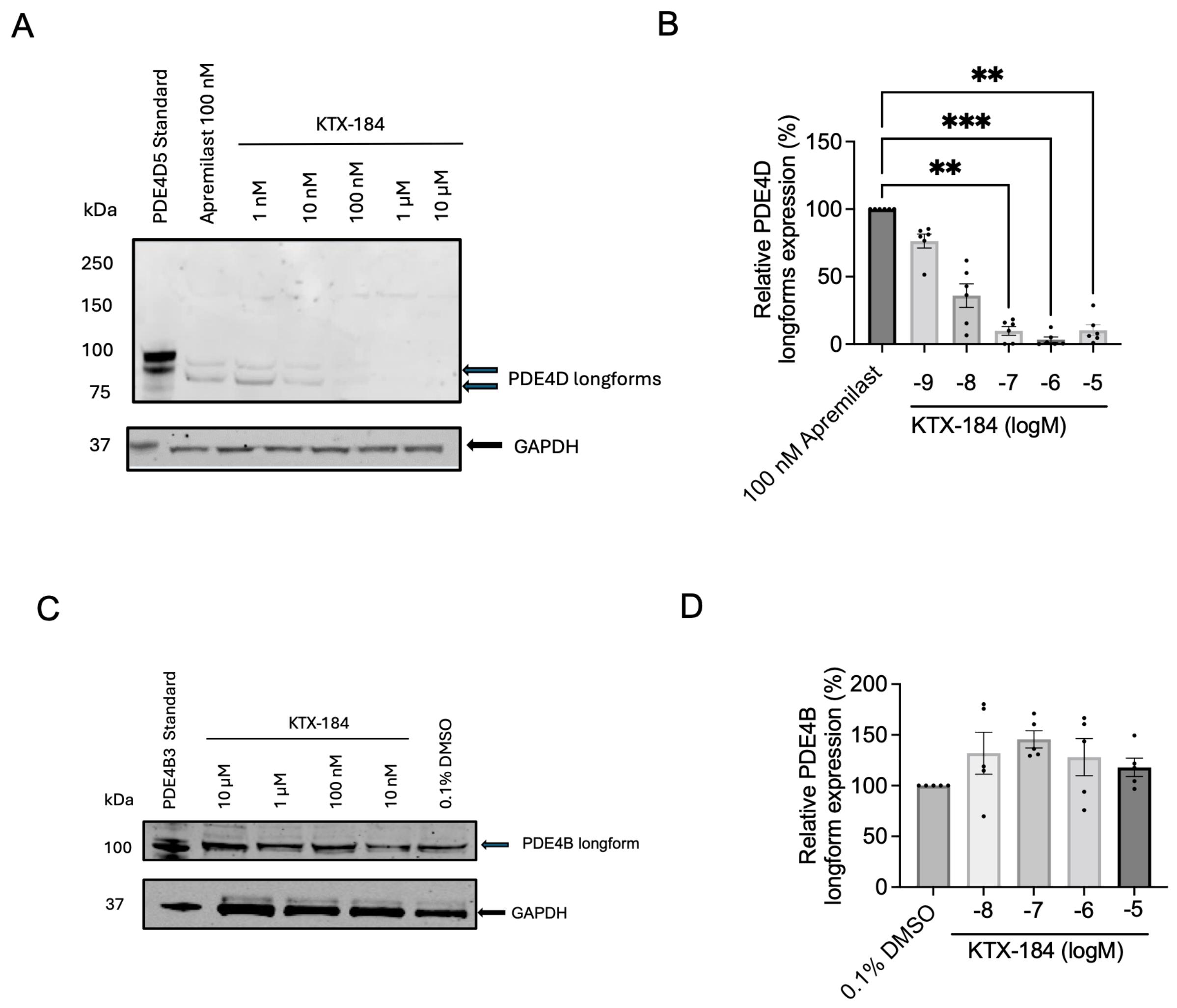


**Supplementary Figure 1. KTX-184 concentration-dependently reduces PDE4D long-form protein abundance without significantly altering PDE4B long-form expression H9c2 cardiomyoblasts.**

**a,** Representative Western blot showing PDE4D long-form expression following treatment with increasing concentrations of KTX-184 (1 nmol/L–10 µmol/L). Recombinant PDE4D5 standard and apremilast (100nmol/L) were included as controls. GAPDH was used as a loading control. **b,** Densitometric quantification of PDE4D long-form expression following treatment with KTX-184 (1 nM–10 µmol/L), expressed relative to apremilast (100nmol/L) control (100%). **c,** Representative Western blot showing PDE4B long-form expression following treatment with KTX-184 (10 nmol/L–10 µmol/L). Recombinant PDE4B3 standard and vehicle (0.1% DMSO) were included as controls. GAPDH was used as a loading control. **d,** Densitometric qquantification of PDE4B long-form expression following treatment with KTX-184 (1 nmol/L–10 µmol/L), expressed relative to the 0.1% DMSO control (100%) condition. Data presented as mean ± S. E. M., $n = 5$, $^{**}P \leq 0.01$ and $^{***}P \leq 0.001$ based on Kruskal-Wallis test with Dunn's multiple comparison.

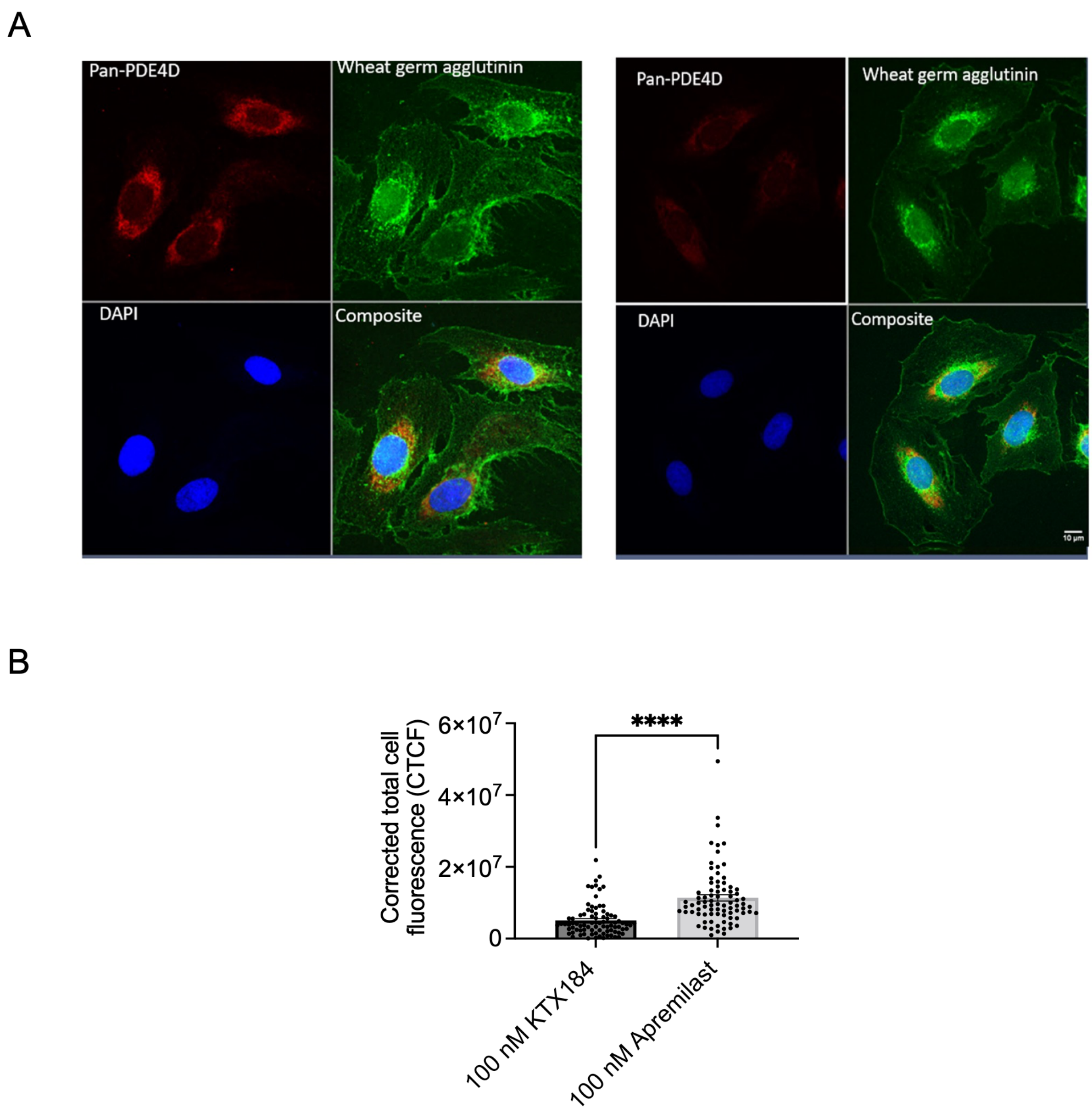


**Supplementary Figure 2. KTX-184 reduces cellular PDE4D abundance compared with apremilast in H9c2 cardiomyoblasts.**

**a,** Representative immunofluorescence images of H9c2 cardiomyoblasts treated with KTX-184 (100 nmol/L) or apremilast (100 nmol/L) and stained for pan-PDE4D (red) and wheat germ agglutinin (WGA; green), with nuclei counterstained with DAPI (blue). Composite images show the merged fluorescence channels. Scale bar, 10 µm. **b**, Quantification of pan-PDE4D immunofluorescence expressed as a corrected total corrected total cell fluorescence

(CTCF) in KTX-184- and apremilast-treated cells. Individual data points are shown, with bars representing mean ± SEM. Data presented as mean mean ± S. E. M., *n* = 82-84, from N = 5 biological replicates , ****$P \leq 0.0001$ based on Welch's *t*-test.

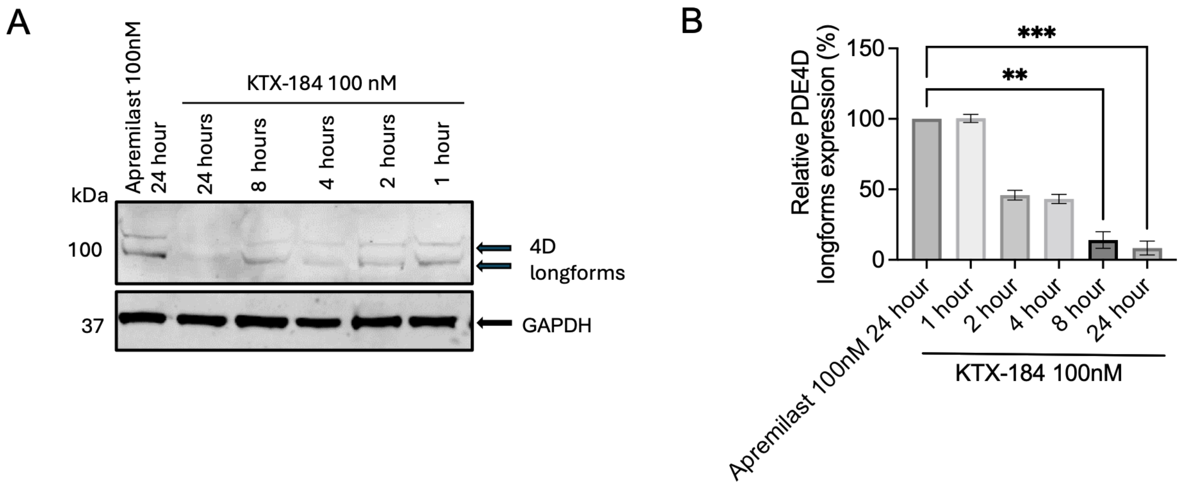


**Supplementary Figure 3. KTX-184 induces a time-dependent reduction in PDE4D long-form protein abundance.**

**a,** Representative Western blot showing PDE4D long-form protein abundance at the indicated time points (1-h – 24-h) following treatment with KTX-184. PDE4D longforms are indicated by arrows, and GAPDH was used as a loading control. **b,** Quantification of relative PDE4D longform expression at the indicated time points following KTX-184 treatment, normalised to the control condition (100%). Data presented as Mean +/- SEM, $n$ = 5, **P ≤ 0.01, ***P ≤ 0.001 based on Kruskal-Wallis with Dunn's multiple comparison.

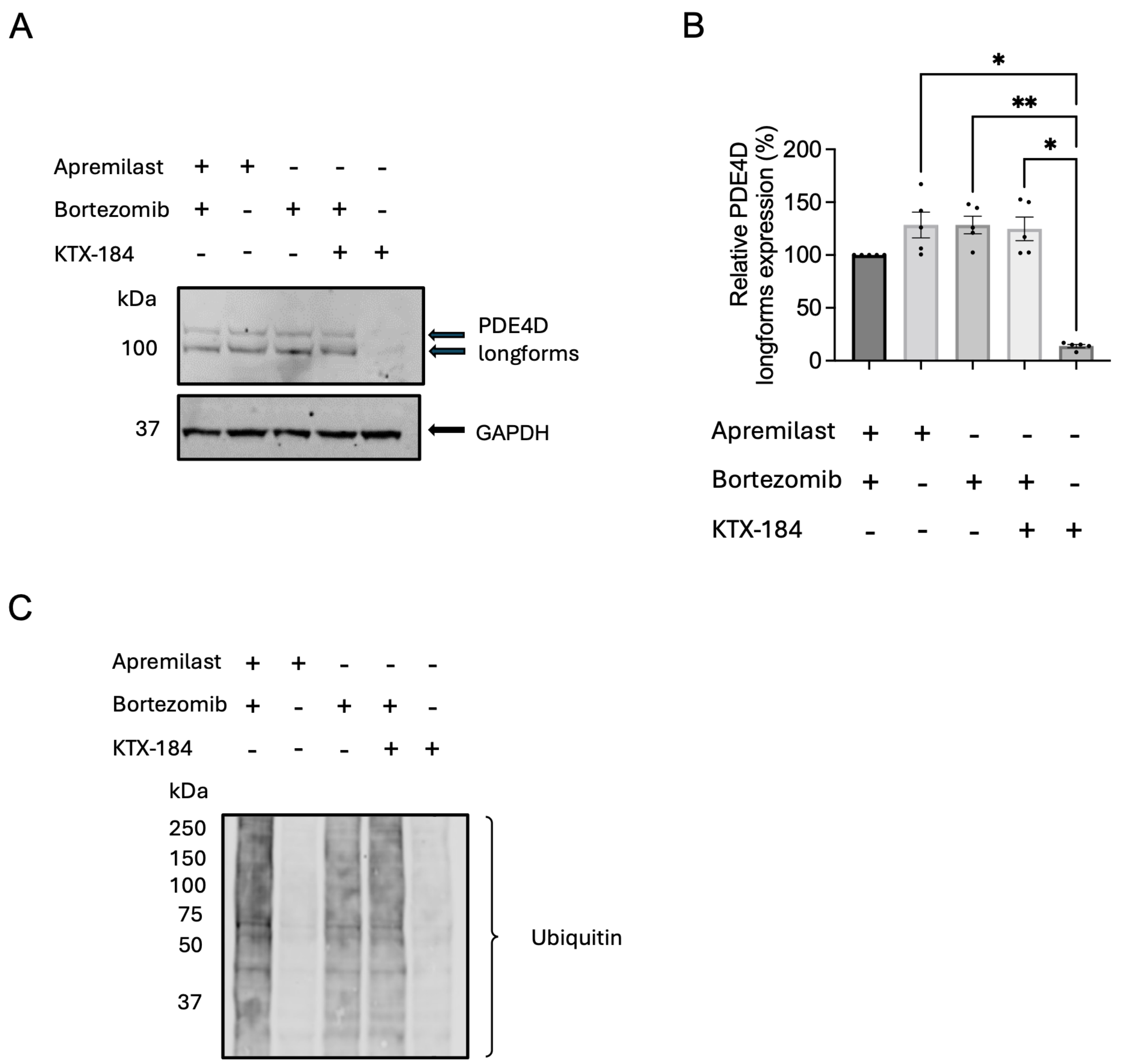


**Supplementary Figure 4. Proteasome inhibition prevents KTX-184-mediated degradation of PDE4D long-forms.**

**a**, Representative Western blot showing PDE4D long-form protein abundance following treatment with KTX-184 or apremilast in the presence or absence of the proteasome inhibitor bortezomib. PDE4D long-forms are indicated by arrows, and GAPDH was used as a loading control. **b**, Densitometric quantification of PDE4D long-form protein abundance under the indicated treatment conditions, normalised to the control conditions. **c,** Representative Western blot of ubiquitinated protein following treatment with KTX-184 or apremilast in the presence or absence bortezomib. Data mean ± S. E. M. $n$ = 5, *$P \leq$ 0.05 and **$P \leq$ 0.01 based on the Kruskal-Wallis test with Dunn's multiple comparison.

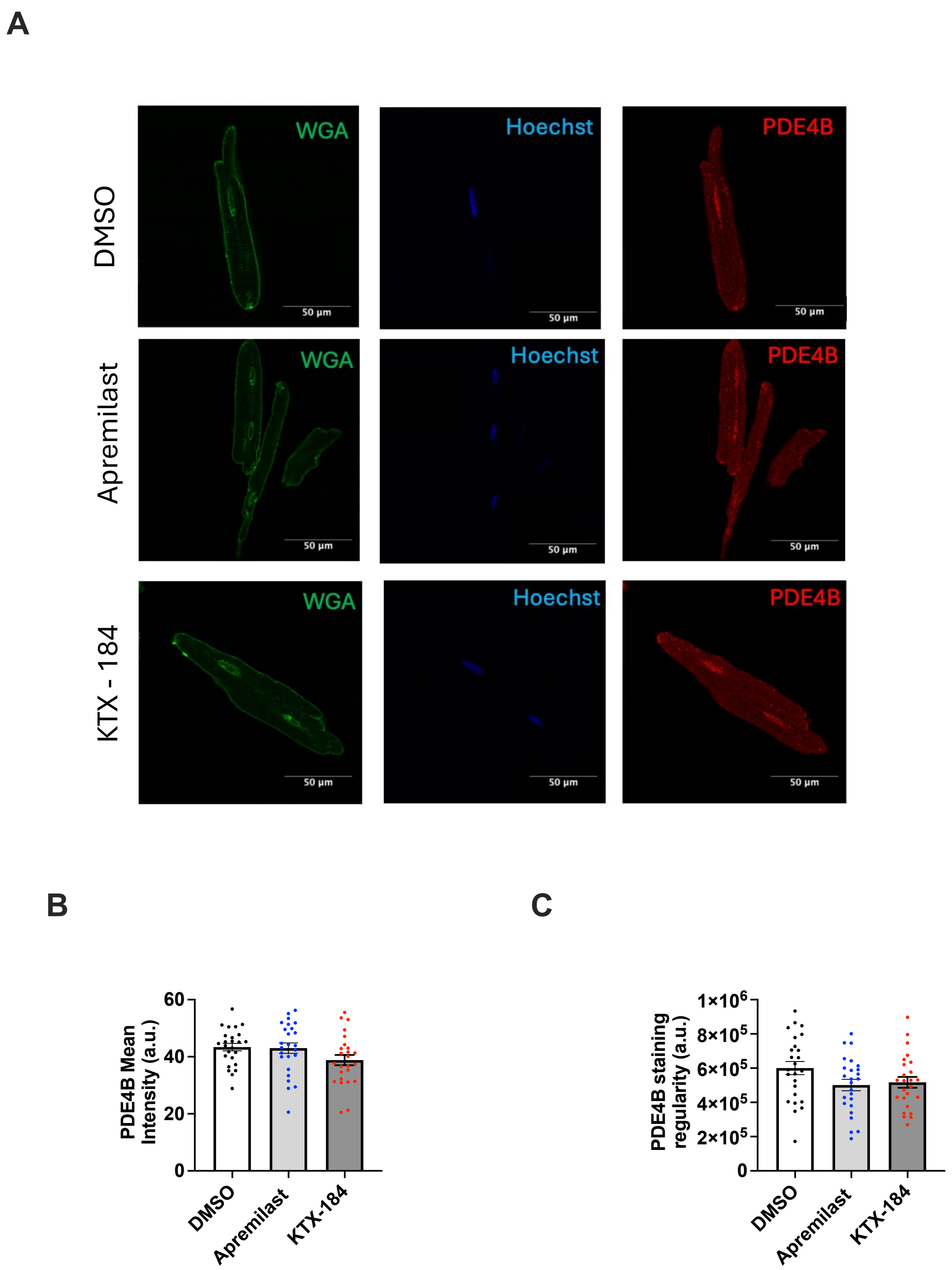


**Supplementary Figure 5. KTX-184 does not alter PDE4B abundance or staining regularity in adult rat left ventricular cardiomyocytes.**

**a,** Representative confocal images of adult rat left ventricular (LV) cardiomyocytes following 24-h treatment with vehicle (0.1% DMSO) apremilast (1 nmol/L) or KTX-184 (1 nmol/L) . Cardiomyocytes were labelled for PDE4B (red) and wheat germ agglutinin (WGA; green), with nuclei counterstained with Hoechst (blue). Scale bars, 50 µm. **b, c,** Quantification of mean PDE4B fluorescence intensity (**B**) and PDE4B staining regularity (**C**) under the indicated treatment conditions.

Individual points represent cardiomyocytes, with colours denoting treatment groups. Data are presented as mean ± S.E.M, *n* = 25 from 5 rats, no statistically significant differences were detected between treatment groups based on one-way ANOVA with Tukey's multiple comparison test.

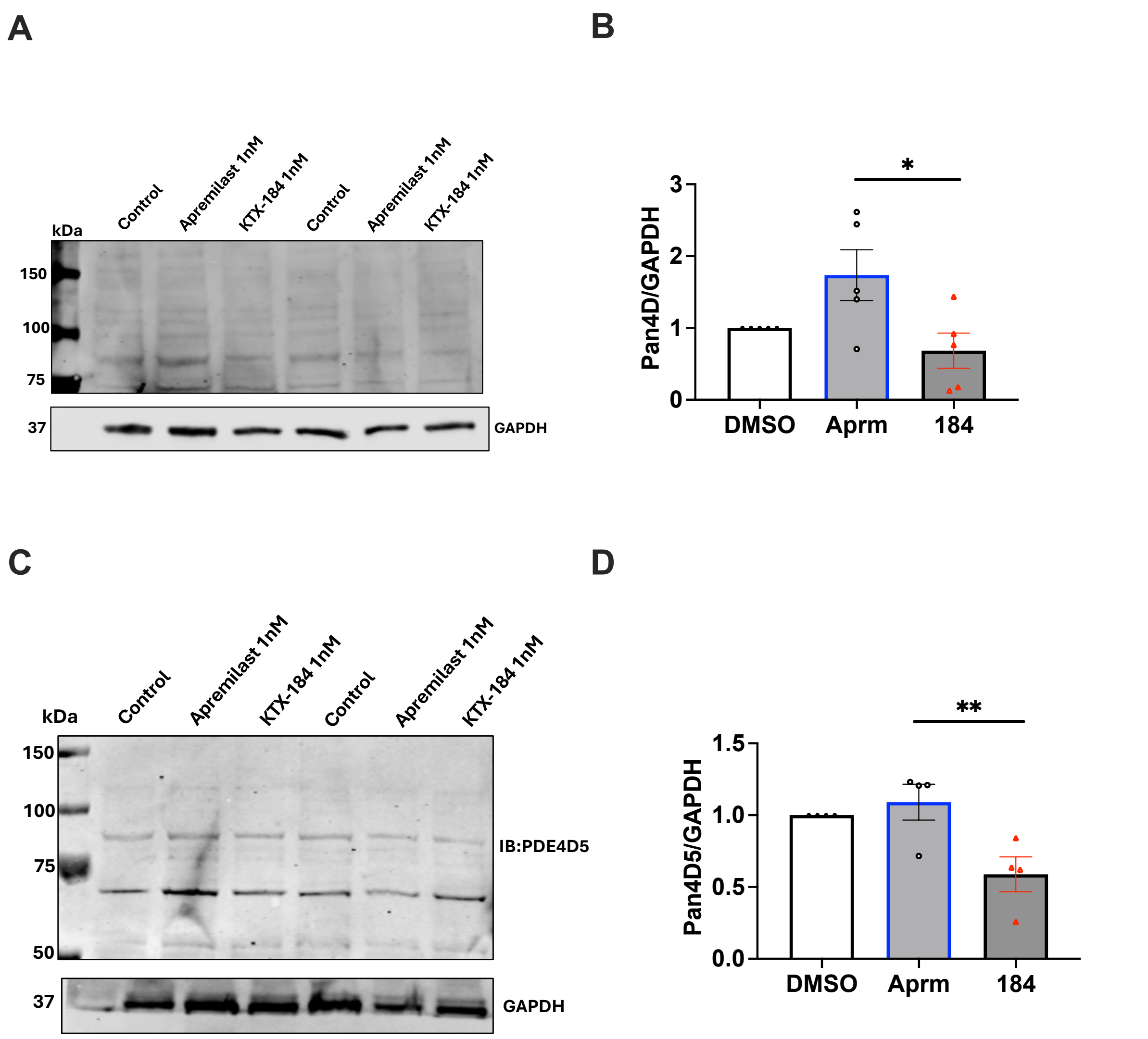


**Supplementary Figure 6**. **KTX-184 reduces PDE4D and PDE4D5 protein abundance in adult rat cardiomyocytes.**

**a**, Representative Western blot showing PDE4D protein abundance in adult rat cardiomyocytes following treatment with vehicle (DMSO), apremilast or KTX-184. GAPDH was used as a loading control. **b**, Densitometric quantification of PDE4D protein abundance under the indicated treatment conditions, normalised to the vehicle control. **c**, Representative Western blot showing PDE4D5 protein abundance in adult rat cardiomyocytes following treatment with indicated conditions. GAPDH was used as a loading control. **d**, Densitometric quantification of PDE4D5 protein abundance under the indicated treatment conditions, normalised to the vehicle control. Data presented as mean ± SEM $n$ = 4-5, *$P \leq 0.05$, ** $P \leq 0.01$ based on the one-way ANOVA with Tukey's multiple comparison test.

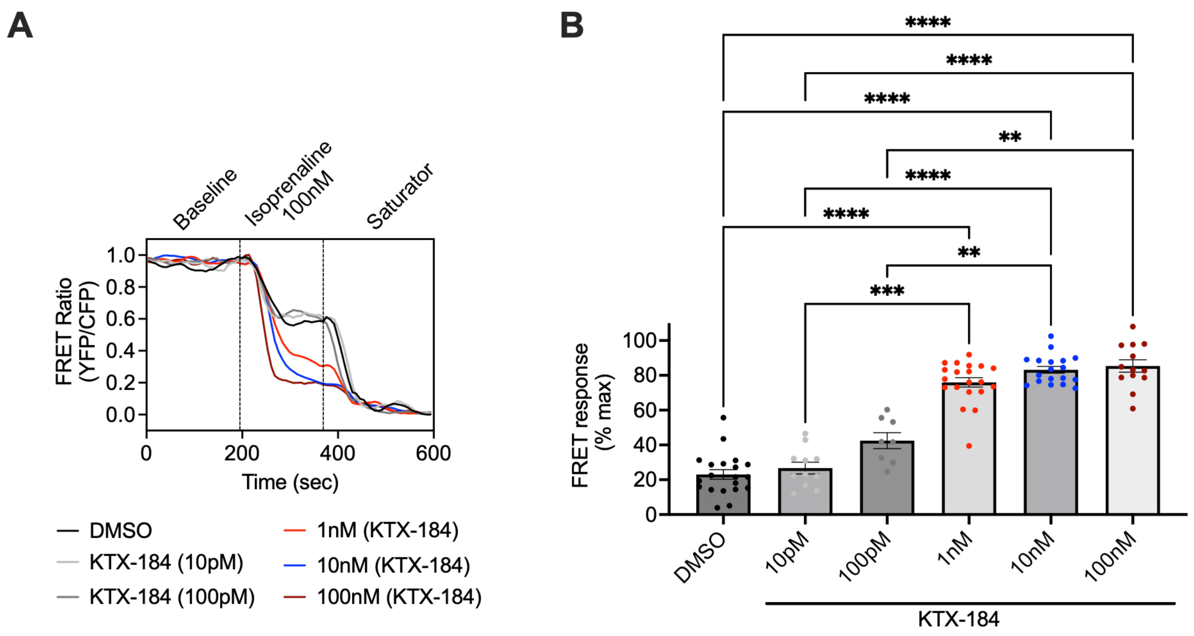


**Supplementary Figure 7. KTX-184 concentration-dependently enhances global cAMP responses in adult rat left ventricular cardiomyocytes.**

**a**, Representative normalised Epac1-camps FRET ratio traces from adult rat left ventricular (LV) cardiomyocytes following 24-h treatment with the indicated concentrations of KTX-184 and stimulation with isoprenaline (100 nmol/L) and saturating agents (the 3-isobutyl-1-methylxanthine [IBMX; 100 µmol/L] plus forskolin [10 µmol/L]. **b**, Quantification of the normalised Epac1-camps FRET response following isoprenaline stimulation in cardiomyocytes treated with the indicated concentrations of KTX-184. Data presented as mean ± S. E. M., $n$ = 8 - 23, $^{*}P \leq 0.05$ $^{**}P \leq 0.01$, $^{***}P \leq 0.001$, $^{****}P \leq 0.0001$ based on the mixed-effect analysis with Dunnett's multiple comparison.

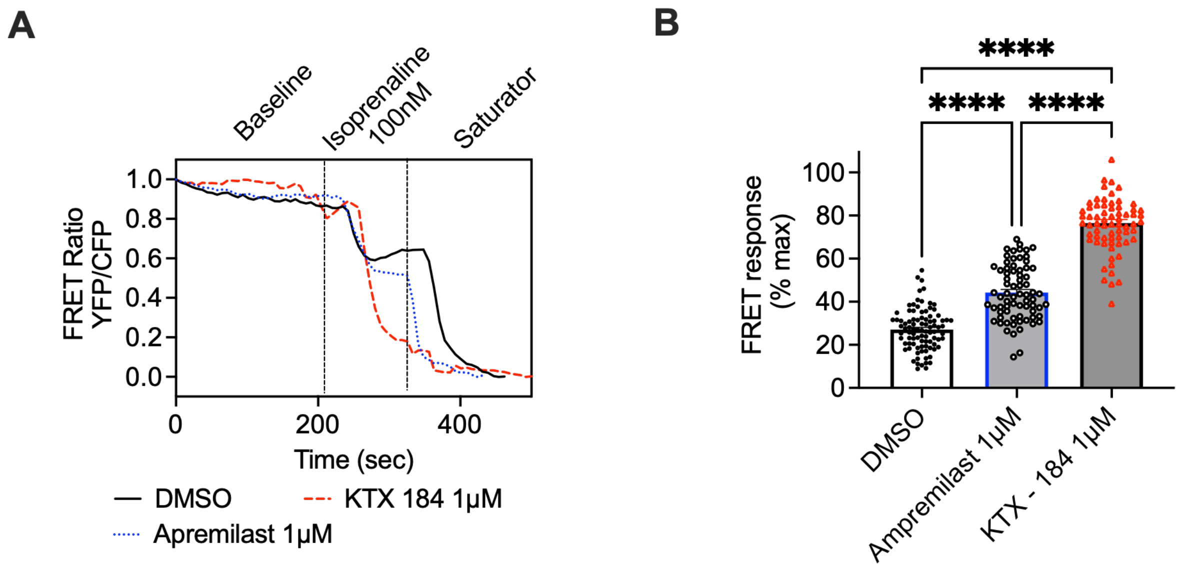


**Supplementary Figure 8. KTX-184 and apremilast enhance global cAMP responses in adult rat left ventricular cardiomyocytes at micromolar concentrations.**

**a,** Representative normalized Epac1-camps FRET ratio traces from adult rat left ventricular (LV) cardiomyocytes following 24-h treatment with vehicle (DMSO), apremilast (1 µmol/L) or KTX-184 (1 µmol/L) and subsequent isoprenaline (100nmol/L) stimulation, followed by saturating conditions the 3-isobutyl-1-methylxanthine (IBMX; 100 µmol/L) plus forskolin (10 µmol/L). **b,** Quantification of the normalised Epac1-camps FRET response under the indicated treatment conditions. Data mean ± S. E. M. $n$ = 69 – 92 from 5 animals, ****$P \leq$ 0.0001 based on the one-way ANOVA with Tukey's multiple comparison.

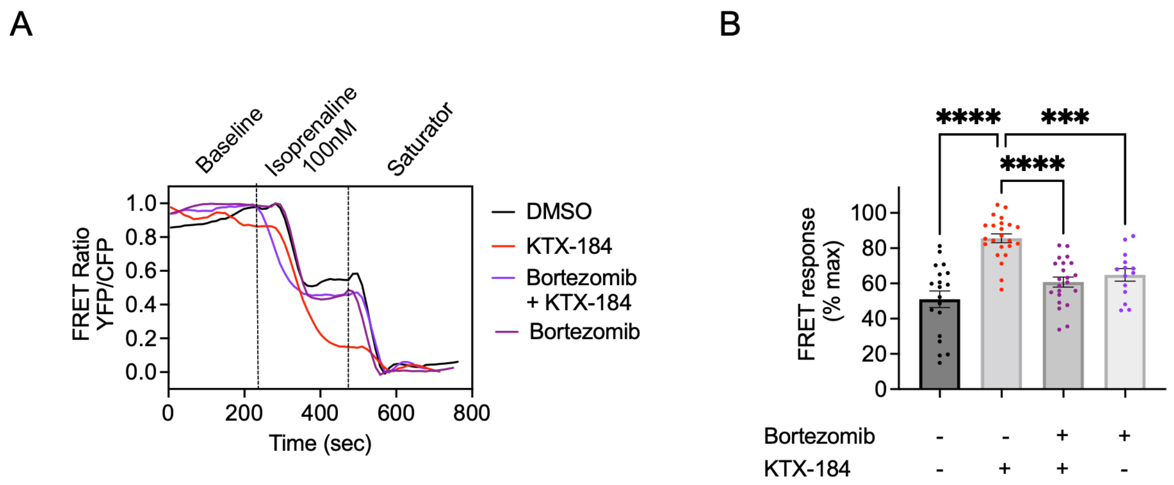


**Supplementary Figure 9. Proteasome inhibition prevents KTX-184-mediated enhancement of global cAMP responses in adult rat left ventricular cardiomyocytes.**

**a**, Representative normalised Epac1-camps FRET ratio traces from adult rat left ventricular (LV) cardiomyocytes following stimulation with KTX-184 (1nmol/L) in the presence or absence of the proteasome inhibitor bortezomib (100 nmol/L) and subsequent stimulation with isoprenaline (100 nmol/L). **b,** Quantification of the normalized Epac1-camps FRET response following isoprenaline and saturating agonists the 3-isobutyl-1-methylxanthine (IBMX; 100 µmol/L) plus forskolin (10 µmol/L) stimulation under the indicated treatment conditions. Data presented as mean ± S. E. M. ***$P \leq$ 0.001, ****$P \leq$ 0.0001, $n$ = 14 - 23 based on based on one-way ANOVA with Tukey's multiple comparison.

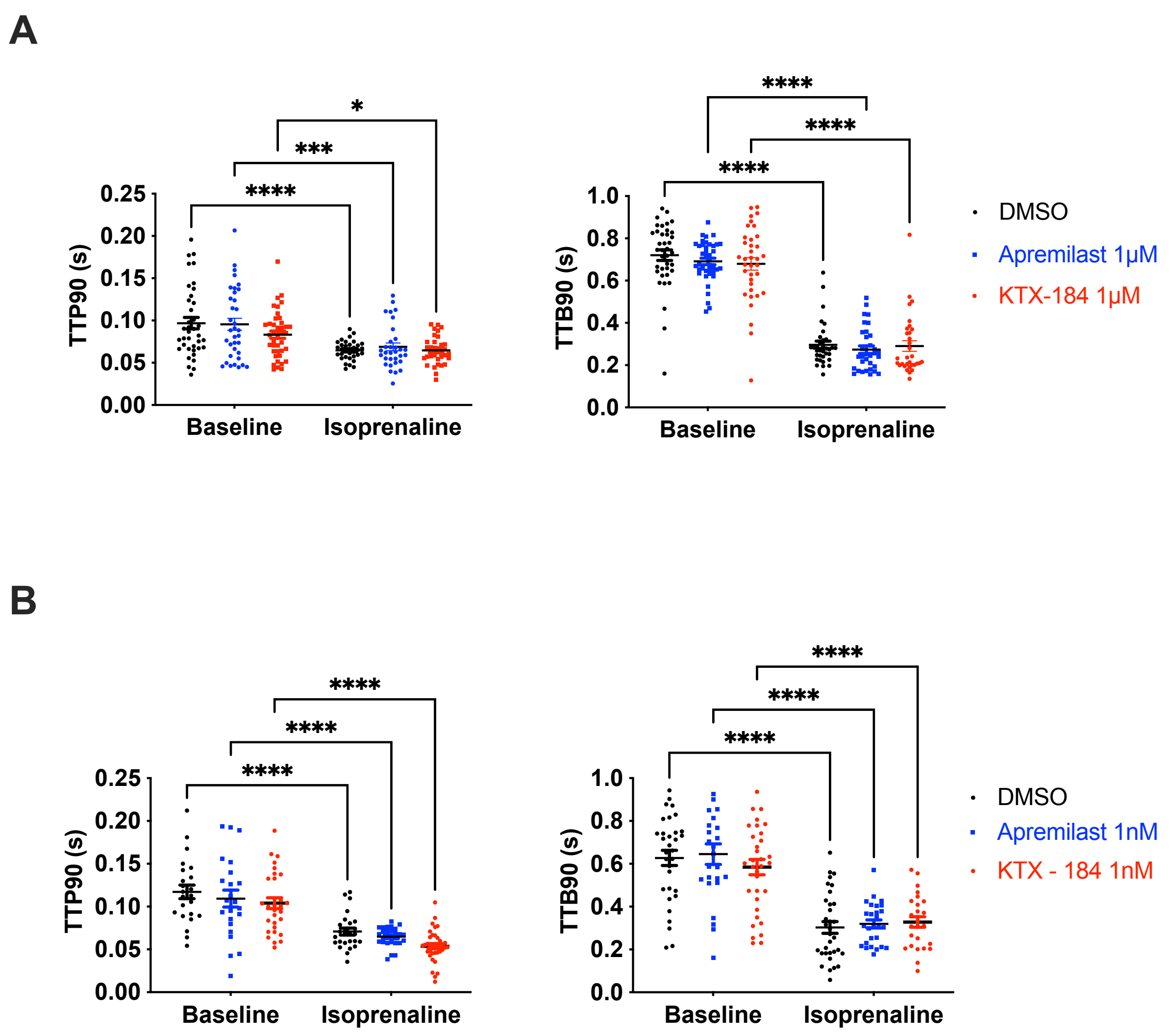


**Supplementary Figure 10. Effect of KTX-184 on contraction and relaxation kinetics in adult rat left ventricular cardiomyocytes.**

**a**, Time to 90% peak contraction (TTP90) and time to 90% return to baseline (TTB90) measured at baseline and following isoprenaline (100 nmol/L) stimulation in adult rat left ventricular (LV) cardiomyocytes treated with vehicle (0.1% DMSO; black), apremilast (1 μmol/L; blue), and KTX-184 (1 μmol/L; red). **b**, TTP90 and TTB90 measured in cardiomyocytes treated with vehicle (0.1% DMSO; black), apremilast (1 nmol/L; blue), and KTX-184 (1 nmol/L; red). Data presented as mean ± S. E. M., *n* = 43-45 from 5 animals. **P* ≤ 0.05, ****P* ≤ 0.001, *****P* ≤ 0.0001 based on the two-way ANOVA with Dunn's multiple comparisons test

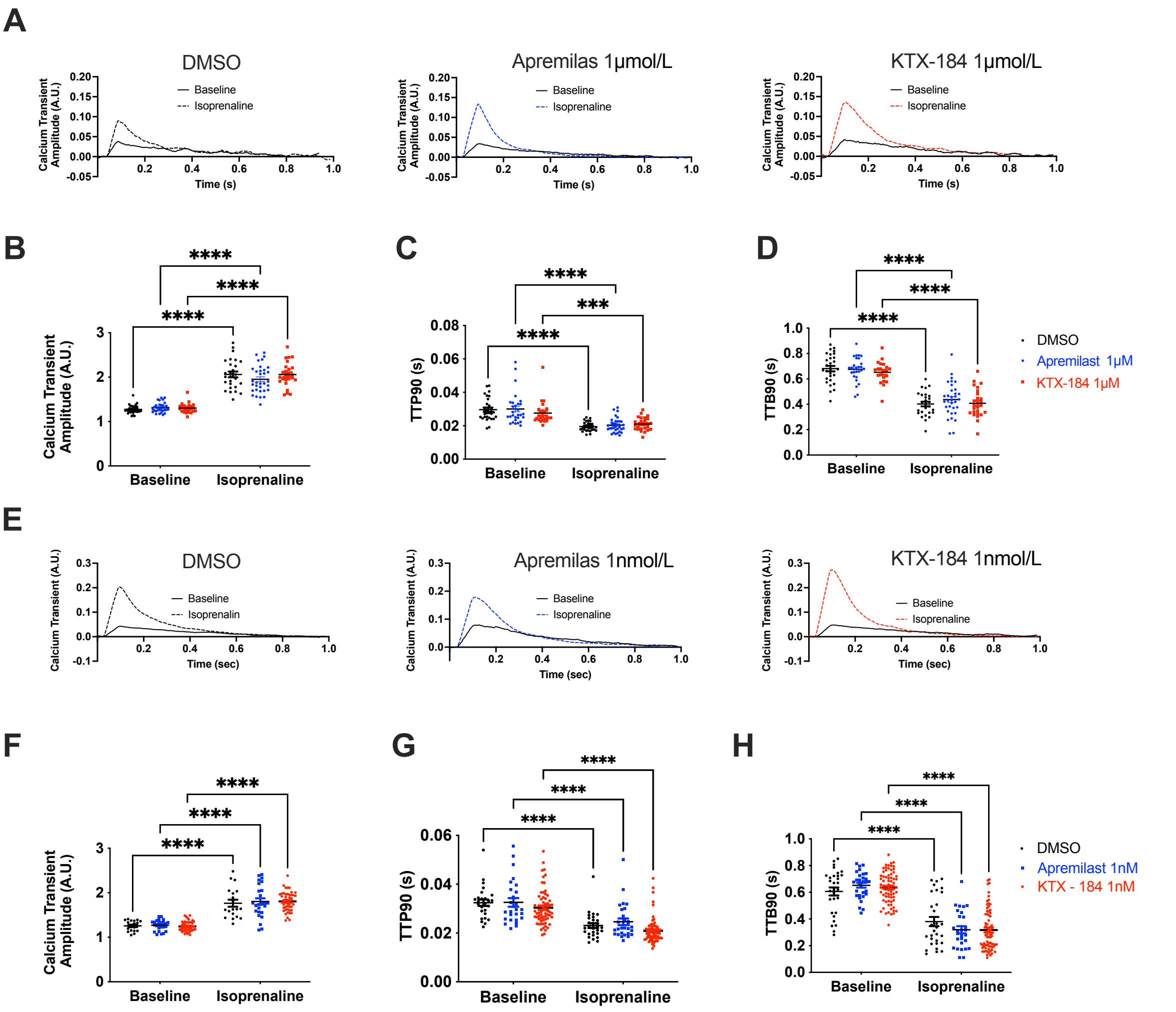


**Supplementary Figure 11. KTX-184 does not alter calcium transient amplitude or kinetics in adult rat left ventricular cardiomyocytes.**

**a**, Representative calcium ($Ca^{2+}$) transients at baseline and following isoprenaline (100nmol/L) stimulation in adult rat left ventricular (LV) cardiomyocytes treated with vehicle (0.1% DMSO); apremilast (1 µmol/L) or KTX-184 (1 µmol/L). **b-d**, Quantification of $Ca^{2+}$ transient amplitude (**B**), time to 90% peak (TTP90; **c)**, and time to 90% return to baseline (TTB90; **d**) at baseline and following isoprenaline (1 µmol/L) stimulation under the indicated treatment conditions. **e**, Representative $Ca^{2+}$ transient amplitude at baseline and following isoprenaline stimulation in cardiomyocytes treated with vehicle (0.1% DMSO), apremilast (1 nmol/L) or KTX-183 (1 nmol/L). **f-h**, Quantification of $Ca^{2+}$ transient amplitude (**f**),TTP90 (**g**), and TTB90 (**h**) at baseline and following isoprenaline (100 nmol/L) stimulation under the indicated treatment conditions. Data presented as mean ± SEM *n* = 43-45 from 5 animals ****$P \leq$ 0.0001 based on the two-way ANOVA with Dunn's multiple comparisons test.

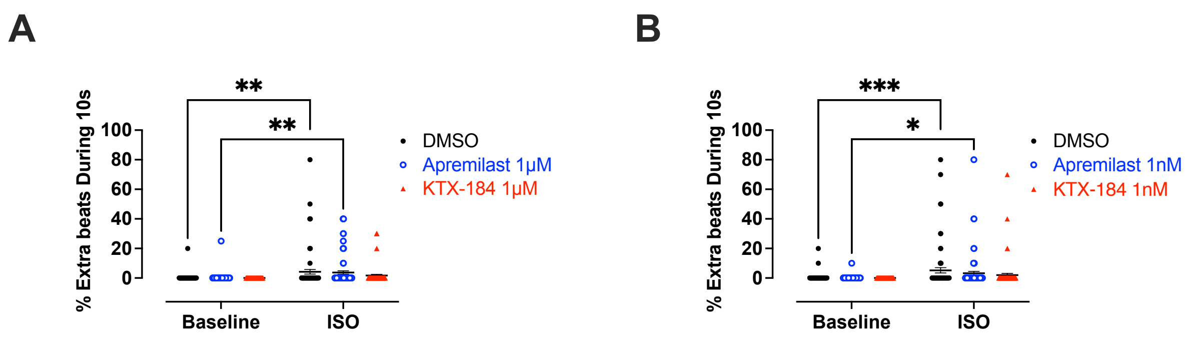


**Supplementary Figure 12. KTX-184 treatment is not associated with a pro-arrhythmic phenotype in adult rat left ventricular cardiomyocytes.**

**a**, Proportion of spontaneous extra beats during a 10-s recording of 1 Hz paced adult left ventricular (LV) cardiomyocytes treated with vehicle (0.1% DMSO), apremilast (1µmol/L) or KTX-184 (1 µmol/L). **b**, Proportion of spontaneous extra beats during a 10-s recording of 1 Hz paced adult LV cardiomyocytes treated with vehicle (0.1% DMSO), apremilast (1nmol/L) or KTX-184 (1nmol/L). Data presented as mean ± S. E. M., $n$ = 73 - 73 from 5 animals. *$P \leq 0.05$, **$P \leq 0.01$, ***$P \leq 0.001$, based on the two-way ANOVA with Šídák's multiple comparisons test

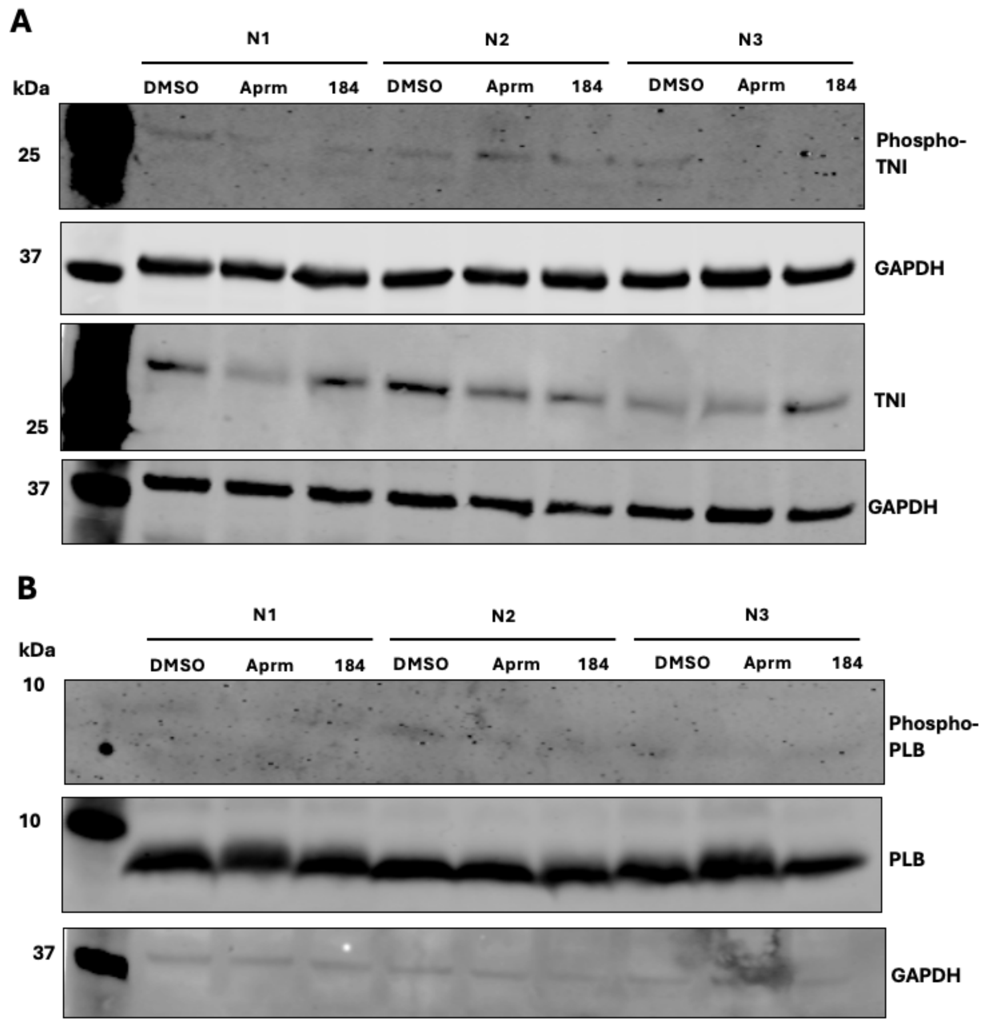


**Supplementary Figure 13. KTX-184 does not alter phosphorylation of troponin I or phospholamban in adult rat left ventricular cardiomyocytes.**

Representative western blots showing phospho-troponin I (Phospho-TNI; a), total troponin I (TNI; **a**), phospho-phospholamban (Phospho-PLB; b), total phospholamban (PLB; **b**), and GAPDH (loading control) in adult rat LV cardiomyocytes following 24-h treatment with vehicle (0.1% DMSO), apremilast (Aprm; 1 nmol/L), or KTX-184 (184; 1 nmol/L). Data are shown from three independent biological replicates (N1–N3). Molecular weight markers are indicated in kDa.

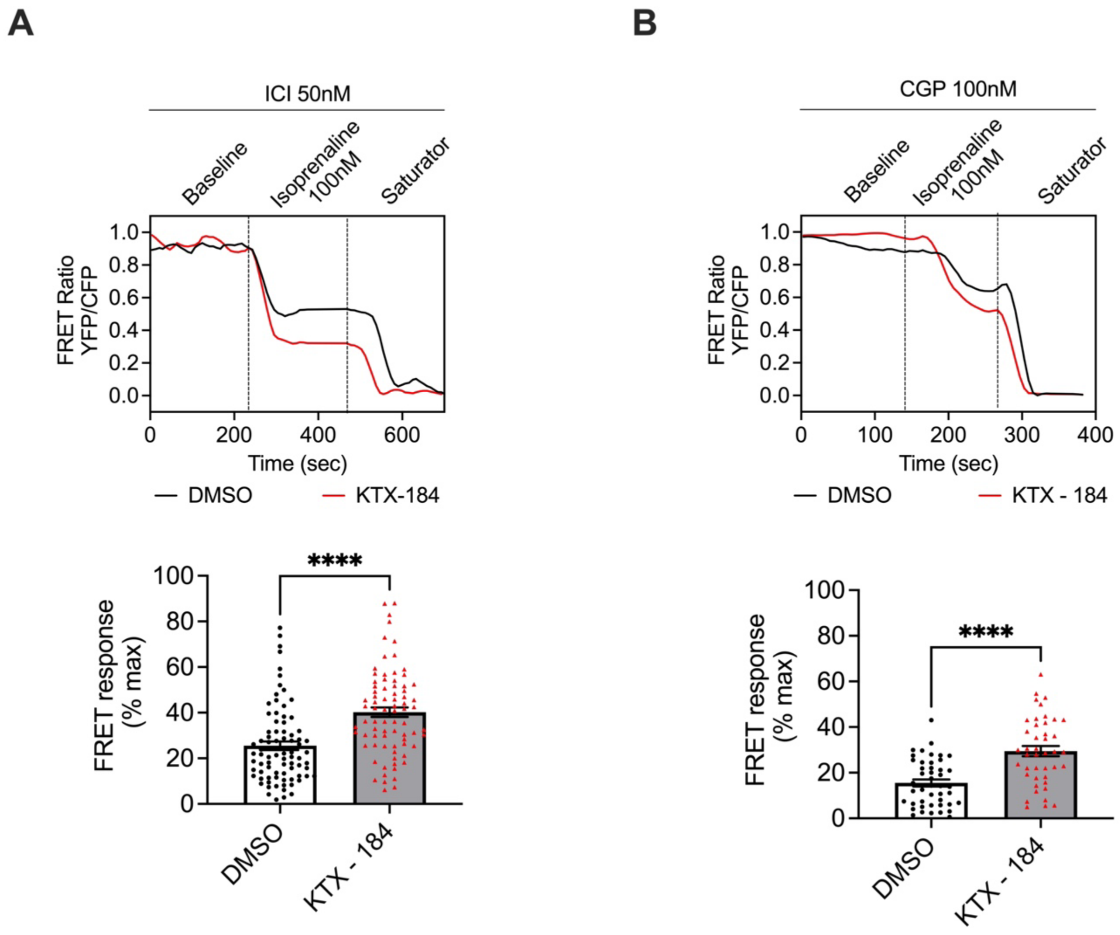


**Supplementary Figure 14. KTX-184 enhances $\beta_1$- and $\beta_2$-adrenergic receptor-mediated global cAMP responses in adult rat left ventricular cardiomyocytes.**

**a,** Representative normalised Epac1-camps FRET ratio trace (top) and quantification of the FRET response (bottom) in adult rat left ventricular (LV) cardiomyocytes treated with vehicle (0.1% DMSO) or KTX-184 in the presence of $\beta_2$-adrenergic receptor antagonist ICI 118,551 (50 nmol/L), followed by stimulation with isoprenaline (100 nmol/L) and saturating agonist conditions the 3-isobutyl-1-methylxanthine (IBMX; 100 µmol/L) plus forskolin (10 µmol/L). **b**, Representative normalised Epac1-camps FRET ratio traces (top) and quantification of the FRET response (bottom) in cardiomyocytes treated with vehicle (0.1% DMSO) or KTX-184 in the presence of the $\beta_1$-adrenergic receptor antagonist CGP 20712A (100 nmol/L), followed by stimulation with isoprenaline (100 nmol/L) and saturating agonist conditions as indicated.

Data presented as mean ± S. E. M.; $n$ = 43-80 from 5 animals $****P \leq 0.0001$ based on *t*-test.

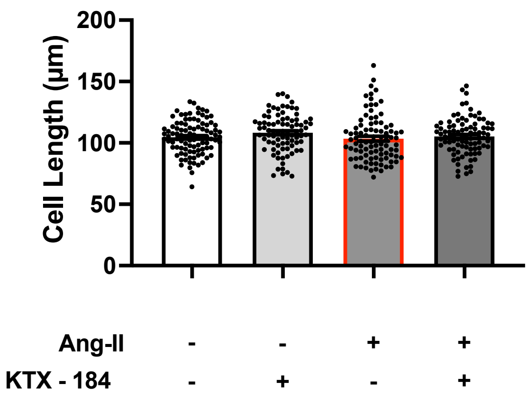


**Supplementary Figure 15. Angiotensin II and KTX-184 do not alter cell length in adult rat left ventricular cardiomyocytes.**

Quantification of cell length in adult rat left ventricular (LV) cardiomyocytes treated with vehicle (0.1% DMSO), KTX-184 (1 nmol/L), angiotensin II (Ang II; 12 µmol/L), or Ang II in combination with KTX-184. Data presented as mean ± S. E. M., *n* = 84 - 97 from 5 animals. No statistically significant differences were detected between treatment groups based on the one-way ANOVA with Tukey's multiple comparison.

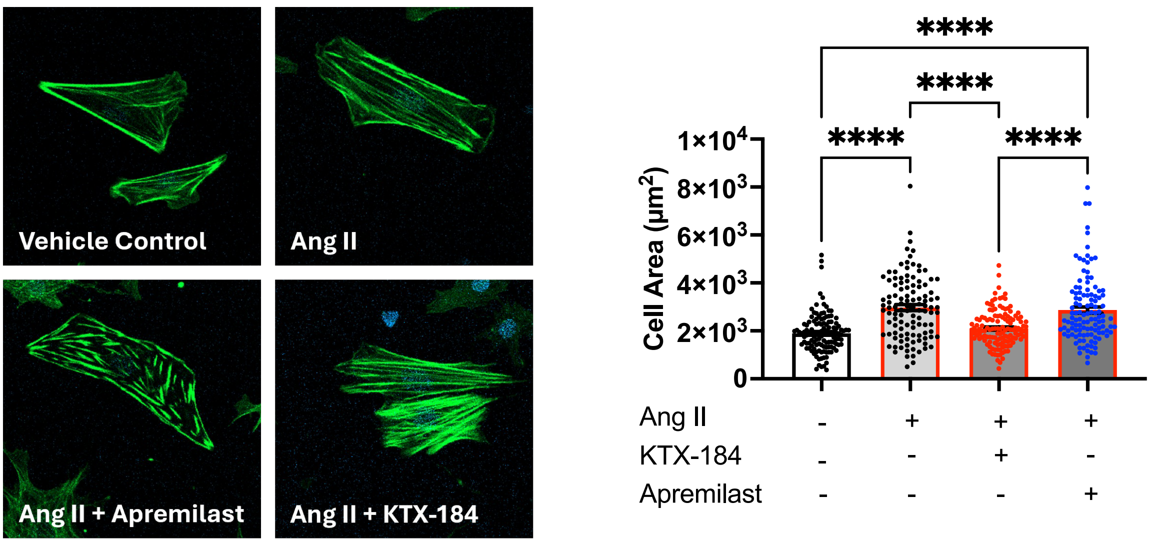


**Supplementary Figure 16. KTX-184 attenuates angiotensin II-induced hypertrophy in neonatal rat cardiomyocytes.**

Left: Representative fluorescence images of neonatal rat cardiomyocytes treated with vehicle (0.1% DMSO), angiotensin II (Ang II; 500nmol/L), Ang II in combination with apremilast (100nmol/L), or Ang II in combination with KTX-184 (100 nmol/L). 24-h before the addition of Ang II at 500 nmol/L. Cardiomyocytes were pre-treated with apremilast or KTX-184 for 24-h before addition of Ang II and incubated for a further 48-h. Cells were fixed and stained with phalloidin-FITC to visualize F-actin. Right: Following a 48h incubation, cells were fixed, stained with phalloidin-FITC, and the cell area was measured. Left - representative images of each treatment group. Right: Quantification of cardiomyocyte cell area under the indicated treatment conditions. Data presented as mean ± S. E. M. from 3 independent experiments, $****P \leq 0.0001$ based on one-way ANOVA, with Dunnett's multiple comparisons test.

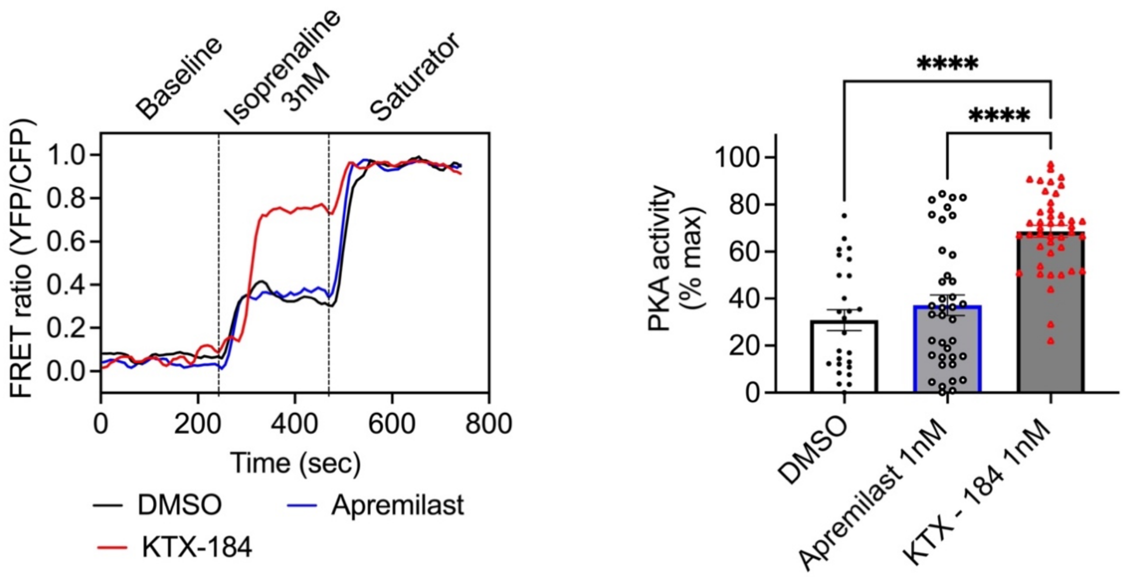


**Supplementary Figure 17. KTX-184 enhances nuclear PKA activity in adult rat left ventricular cardiomyocytes.**

Left: Representative normalised AKAR3-NES FRET ratio traces from adult rat left ventricular (LV) cardiomyocytes following 24-h treatment with vehicle (0.1% DMSO), apremilast (1 nmol/L) or KTX-184 (1 nmol/L), with isoprenaline (3 nmol/L) and saturating agonist (3-isobutyl-1-methylxanthine (IBMX; 100 µmol/L) plus forskolin (10 µmol/L)) conditions. Right: Quantification of the normalized AKAR3-NLS FRET response, reflecting nuclear PKA activity, under the indicated treatment conditions. Data presented as mean ± S. E. M., $n$ = 26 - 43 from 5 animals, ****$P \leq$ 0.0001 based on the one-way ANOVA with Tukey's multiple comparison.

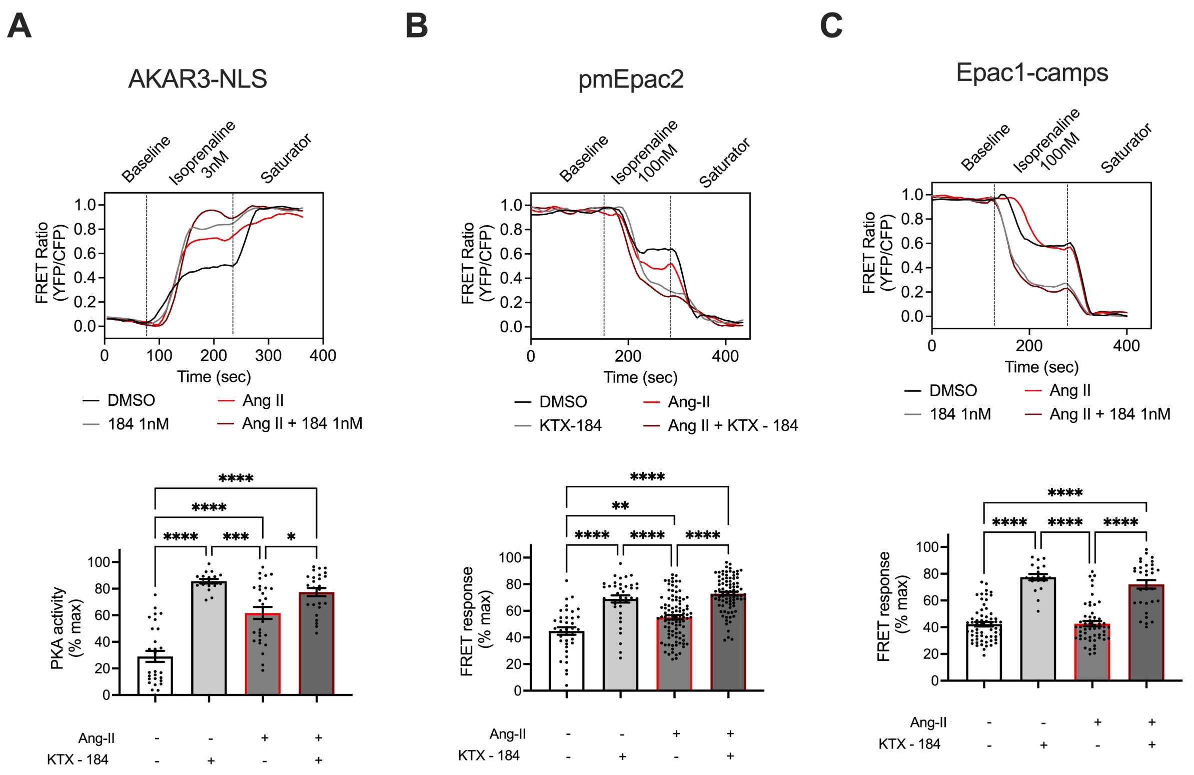


**Supplementary Figure 18. KTX-18 enhances membrane cAMP and nuclear PKA response in angiotensin II-treated adult rat left ventricular cardiomyocytes**.

**a,** Representative normalised AKAR3-NLS FRET ratio traces (top) and quantification of nuclear PKA activity (bottom) in adult rat left ventricular (LV) cardiomyocytes treated with vehicle (0.1% DMSO), KTX-184 (1nmol/L), angiotensin II (Ang II; 12 µmol/L), or Ang II in combination with KTX-184 for 24-h. Cardiomyocytes were stimulation with isoprenaline (3 nmol/L), followed by saturating agent (3-isobutyl-1-methylxanthine (IBMX; 100 µmol/L) plus forskolin (10 µmol/L)). **b**, Representative normalized pmEpac2 FRET ratio traces (top) and quantification of the FRET response (bottom), reflecting membrane-associated cAMP responses, under the indicated treatment conditions. Cardiomyocytes were stimulation with isoprenaline (100 nmol/L), followed by saturating.  **c**, Representative normalized Epac1-camps FRET ratio traces (top) and quantification of the FRET response (bottom), reflecting global cytosolic cAMP responses, under the indicated treatment conditions. Data presented as mean ± S. E. M, $n$ = 17 - 89 from 3 - 5 animals. *$P \leq 0.05$, **$P \leq 0.01$, ***$P \leq 0.001$, ****$P<0.0001$ based on one-way ANOVA with Tukey's multiple comparison.

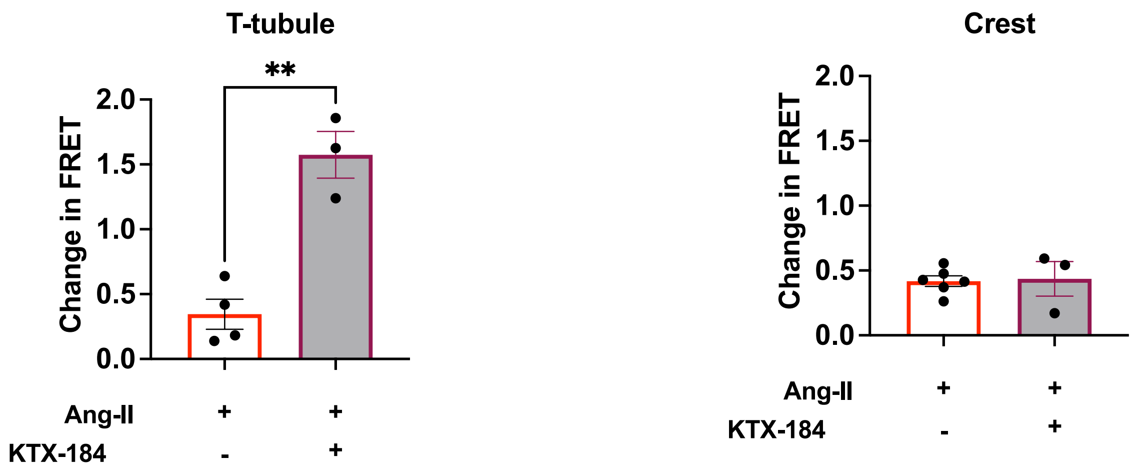


**Supplementary Figure 19. KTX-184 enhances T-tubule-associated, but not crest-associated, membrane cAMP response in angiotensin II-treated adult left ventricular cardiomyocytes.** Quantification of membrane cAMP responses measured using pmEpac cAMP sensor in adult rat left ventricular (LV) cardiomyocytes treated with angiotensin II (Ang II; 12 µmol/L) alone or in combination with KTX-184 (1 nmol/L) for 24-h. Cardiomyocytes were acutely stimulated at T-tubule (left) or crest (right) location with isoprenaline (10 µmol/L. Data presented as mean ± S. E. M.; *n* = 3 – 6 from 3 animals. **P < 0.01, unpaired two-tailed Student’s *t*-test.

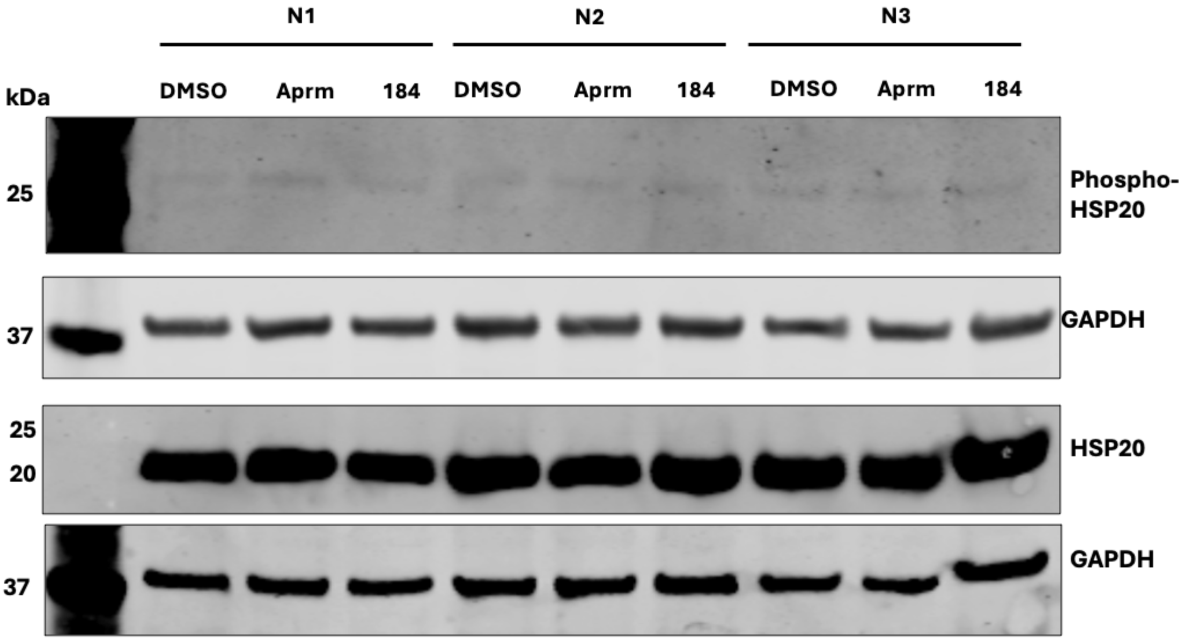


**Supplementary Figure 20. KTX-184 does not alter phosphorylation of HSP20 in adult rat left ventricular cardiomyocytes.**

Representative western blots showing phospho-HSP20 (Phospho-HSP20), total HSP20, and GAPDH (loading control) in adult rat LV cardiomyocytes following 24-h treatment with vehicle (0.1% DMSO), apremilast (Aprm; 1 nmol/L), or KTX-184 (184; 1 nmol/L). Data are shown from three independent biological replicates (N1–N3). Molecular weight markers are indicated in kDa.

**Supplementary Table 1. List of primers for RT-qPCR for mRNA.**

| Gene name | Species | Forward primer | Reverse primer |
|---|---|---|---|
| *Bak1* | rat | GCAGTCAGGCAGGTGACAA | CTCTGTGTCCTGGGCAACTT |
| *Bid* | rat | AGCACGAGGAAAGGAAGGTG | CAGGGAACCACCAGCAGAC |
| *Myh7* | rat | CCGAGTCCCAGGTCAACAAG | TACTCTTCATTCAGGCCCTTGG |
| *Nppa* | rat | GGGGGTAGGATTGACAGGAT | CTCCAGGAGGGTATTCACCA |
| *Nppb* | rat | GAACAATCCACGATGCAGAAGC | TCTGCCCAAAGCAGCTTGAA |

**Supplementary Table 2. List of primers for RT-qPCR for miRNA.**

| miRNA | Species | Mature miRNA sequence | Assay ID |
|---|---|---|---|
| hsa-miR-1 | rat | UGGAAUGUAAAGAAGUAUGUAU | 002063 |
| U6 | rat | GTGCTCGCTTCGGCAGCACATATACTAAAAT TGGAACGATACAGAGAAGATTAGCATGGC CCCTGCGCAAGGATGACACGCAAATTCGTG AAGCGTTCCATATTTT | 001973 |
